# Blood vessels guide Schwann cell migration in the adult demyelinated CNS through Eph/ephrin signaling

**DOI:** 10.1101/498261

**Authors:** Beatriz Garcia-Diaz, Corinne Bachelin, Fanny Coulpier, Gaspard Gerschenfeld, Cyrille Deboux, Violetta Zujovic, Patrick Charnay, Piotr Topilko, Anne Baron-Van Evercooren

## Abstract

Schwann cells (SC) enter the central nervous system (CNS) in pathophysiological conditions. However, how SC invade the CNS to remyelinate central axons remains undetermined. We studied SC migratory behavior *ex vivo* and *in vivo* after exogenous transplantation in the demyelinated spinal cord. Data highlight for the first time that SC migrate preferentially along blood vessel in perivascular extracellular matrix (ECM), avoiding CNS myelin. We demonstrate *in vitro* and *in vivo* that this migration route occurs by virtue of a dual mode of action of Eph/ephrin receptor. Indeed, EphrinB3, enriched in myelin, interacts with SC Eph receptors, to drive SC away from CNS myelin, and triggers their preferential adhesion to ECM components, such as fibronectin via integrinβ1 interactions. This complex interplay enhances SC migration along the blood vessel network and together with lesion-induced vascular remodeling facilitates their timely invasion of the lesion site. These novel findings elucidate the mechanism by which SC invade and contribute to spinal cord repair.

## INTRODUCTION

Myelination, the evolutionary characteristic acquired by vertebrates to allow rapid and saltatory nerve conduction, is supported by two different glial cell types, oligodendrocytes in the central nervous system (CNS), and Schwann cells (SC) in the peripheral nervous system (PNS). These two cell types are mutually exclusive in physiological conditions. However, in demyelinating diseases or injury, this PNS/CNS segregation is compromised and both, SC can invade and repair the CNS(1, 2), and oligodendrocytes myelinate peripheral nerve root axons(3). Remyelination of CNS axons by SC protects axons, restores axonal conduction and even reverses neurological deficits(4), highlighting their potential to rescue the injured CNS(5). These remyelinating SC arise either from the PNS(6, 7), or are generated from adult oligodendrocyte precursors cells (OPC)(8, 9). However, SC remyelination of CNS axons is always restricted to the spinal roots entry and exit zones (reviewed in(5)), and the presence of peripheral myelin has been frequently observed close to blood vessels (BV)(1). These observations suggest that although SC migrate efficiently *in vitro*(10, 11) and *in vivo*(12) into the PNS, their survival and migration within the CNS are limited. The presence of peripheral myelin close to BV raises the possibility for BV to play a role in guiding SC movements within the CNS. While migration along BV has been recently described for different CNS cell types(13-15) including for SC along regenerative nerves(12), their role in SC invasion of the CNS has not been explored.

Astrocytes(16, 17) and CNS white matter(18-20) inhibit SC migration within CNS, pointing out that multiple CNS components can alter this process. Among the molecules involved in various cell type segregation and guidance within the CNS, the Eph/ephrin family has been implicated in both developmental(21, 22) and pathological conditions(23). In particular, EphrinB3, is expressed in myelin in brain and mouse spinal cord(24-26), and plays an important role in preventing neurite outgrowth(24), axonal regeneration(25) and oligodendrocyte progenitor differentiation(26). EphrinB3 acts also as a repellent molecule in the guidance of axon tracts towards the spinal cord during development(27). Interestingly, SC express several Ephrin receptors, including EphA4 that mediates SC repulsion by astrocyte EphrinAs(28). Hence, these receptors could mediate also similar SC repulsion by myelin-associated EphrinB3. In addition, EphB is involved in SC sorting and migration in regenerating peripheral nerves(29). Eph/ephrin signalling is involved in the modulation of cell-cell adhesion, which results in increased integrin-mediated adhesion of Eph/ephrin-expressing cells(30, 31) or, as in the case of EphB signaling, in cell sorting by re-localization of N-cadherin in SC(29). Furthermore, EphrinB ligands control cell migration through positive adhesion to substrates such as collagen and fibronectin (FN)(32, 33), main components of perivascular extracellular matrix (ECM). Therefore, these cues could influence the capacity of SC to migrate within the CNS and/or interact with CNS myelin.

In spite of these observations, a role for BV in SC migration/recruitment in the CNS and for EphrinB3 in modulating SC interactions with CNS myelin and/or BV has not been investigated. Using *ex vivo, in vivo* and *in vitro* paradigms, we show that BV are a preferred substrate for SC migration into the CNS, and that, angiogenesis and BV remodeling constitute a physiological response to demyelination. Moreover, we establish that CNS myelin plays an essential role in SC exclusion from the CNS and demonstrate that this effect is partially mediated by EphrinB3. Finally, myelin-associated EphrinB3 modulates SC adhesion to the ECM component FN, via interactions with integrinβ1. This increased SC-FN adhesion, to perivascular ECMoverrules SC inhibition by myelin and promotes SC migration along BV, facilitating their arrival at the lesion. Thus Eph/ephrin may guide SC within the CNS according to a dual mode, repulsing SC from CNS white matter on one hand, and favoring their interaction with BV on the other. These observations shed new light on the mechanism of SC invasion into the damaged CNS.

## RESULTS

### Schwann cells preferential migration along blood vessels by passes their inhibition by myelin

To address this question, we analysed the SC interactions with CNS components, myelin and blood vessels, in *ex vivo* and *in vivo* paradigms (Fig. 1, Fig. S1 and Fig. S2). For *ex-vivo* analysis, GFP-expressing SC (GFP^+^) were seeded on frozen spinal cord sections. We found that GFP^+^SC adhered 11.88 fold more over grey matter (0.32±0.05% of total area) than white matter (3.8±0.2% of total area) (Fig S1A, C). Moreover, SC spreading over white matter was reduced compared to grey matter (GFP^+^SC area per Hoechst^+^ in GM: 230.0±8.5 μm^2^/cell; in WM: 172.3±11.91 μm^2^/cell) (Fig. S1D). Immunolabeling for Glut1 showed that BV were 7.78 fold more abundant in grey matter than white matter (Glut1^+^ area in GM: 40.9±8.2% of total area; in WM: 5.2±1.6% of total area) (Fig. S1B,E) and that most GFP^+^ SC (>86%) were in contact with collagen 4-positive BV (Fig. S1F), indicating that SC densities in grey and white matter were correlated with BV densities in these regions.

**Figure 1.**
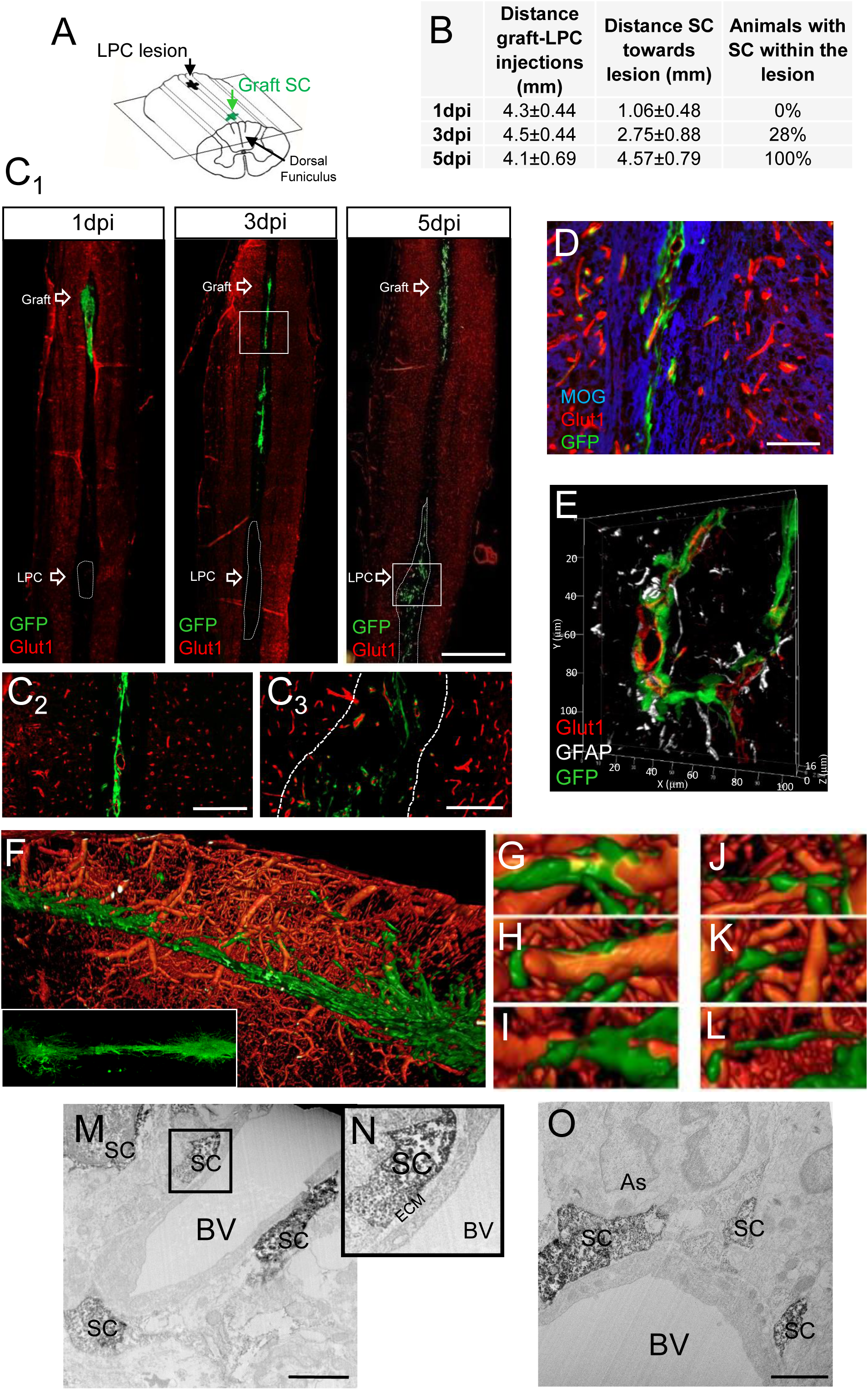
Migrating behavior of SC grafted in white matter remotely from LPC-induced demyelination. Scheme of LPC lesion and SC graft targeted into the dorsal funiculus of the spinal cord; graft and lesion are 4mm apart. (**B**) Extent of migration and percentage of animals with lesions containing SC at different times: SC arrival at the lesion is timely regulated. Data are expressed as mean ± SD, 1dpi (n=5), 3dpi (n=6), 5dpi (n=7). (**C_1_**) General view of longitudinal sections of the spinal cord illustrating the graft and lesion sites at 1dpi, 3dpi and 5dpi, scale bar 1000 µm; (**C_2-_C_3_**) GFP^+^SC grafted within dorsal spinal cord white matter migrate preferentially in close contact with Glut1^+^ endothelial cells when progressing along the midline, towards the lesion (**C_2_**) and spreading within the lesion (**C_3_**), dotted lines identify the lesion sites, scale bar 200 µm. (**D**) Grafted GFP^+^SC along the midline avoid MOG^+^ myelin and are associated with Glut1^+^BV, scale bar 100 µm. (**E**) 3D Z stack reconstruction illustrating GFP^+^SC located between Glut1^+^endothelial cells and GFAP+ perivascular astroglial end feet. (**F-L**) 3D reconstruction after light sheet imaging of clarified whole spinal cord illustrating in (**F**) the abundant vascular network and tdTomato^+^SC migrating from the graft along the dorsal funiculus midline en route for the lesion; (**G-I**) most tdTomato^+^SC exiting the graft are polarized on BV or (**J-L**) between BV evoking jumping events. (**M-O**) Immuno-EM of GFP^+^SC in the midline illustrates several GFP^+^SC revealed by DAB embedded in perivascular ECM (**M**) and between BV and astrocytes (As) (**O**), scale bar 2 µm. (**N**) Higher magnification of the boxed area in (**M**).

To explore how endogenous SC invade the CNS in response to demyelination, we used the Krox20Cre driver line crossed over Rosa-YFP to track the SC lineage (3, 34). Krox20 transcription factor is specific of the PNS, and is expressed in the SC lineage until adulthood. LPC injections were performed in the dorsal funiculus of adult mice (2-3month-old), and mice were sacrificed at 3 days post injection (dpi). YFP^+^SC were detected only in the spinal cord cross sections of demyelinated mice (Fig. S2A,B) but not in controls (Fig S2C,D). YFP+ cells were Sox10^+^ (Fig. S2E,F) but never Olig2+ (Fig S2G) validating their PNS glial origin. We next examined the location of the invading SC-derived population and found that at this early time-point, the YFP^+/^Sox10^+^SC within white mater were found exclusively on Glut1^+^BV. This indicates that such as observed *ex-vivo,* endogenous SC preferentially associate with BV when triggered to invade the demyelinating CNS. Although in normal development Krox20 specifically labels the peripheral SC lineage, under pathological conditions Krox20 was also expressed by Iba1+ microglial cells (Fig.S2H,I).

As the Krox20Cre system was limited in tracing a sufficient number of SC invading the CNS from the periphery, we pursued our investigation using an exogenous paradigm in which GFP^+^SC were grafted two vertebrates away from a lysolecithin (LPC) lesion in the dorsal funiculus of adult wild-type mice (Fig 1A). In this paradigm, grafted SC are known to be recruited specifically by the lesion, modeling SC recruitment after CNS injury, whereas they remain at the graft site in the absence of a lesion (20, 35). As our study focused essentially on SC migration/recruitment, grafted animals were sacrificed at early time-points including 1, 3 and 5-day post-LPC injection (dpi) (Fig. 1B). The spatio-temporal distribution of the transplanted GFP-expressing SC was assessed by scanning longitudinal frozen sections of the spinal cord for the GFP signal throughout the sections (20). At 1 dpi, lesion size (0.40 ± 0.06 mm^2^ per section) and GFP^+^SC grafted area (0.16 ± 0.08 mm^2^ per section) showed minor variability among animals validating the lesion-graft paradigm. Analysis of the GFP signal over time confirmed the spatio-temporal progression of SC towards the lesion, which was systematically reached at 5 dpi (Fig. 1B,C).

In the intermediate zone connecting the graft to the lesion, GFP^+^SC migrated preferentially along the midline avoiding myelin (Fig. 1C,D), as previously described after engraftment in shiverer and nude mice(20, 35). In this area, SC were found preferentially in association with Glut1^+^BV forming a narrow stream of cells (Fig. 1C_2_,D). At their arrival in the lesion, GFP^+^SC were no longer confined to that narrow path, but randomly spread within the lesion. Co-detection of GFP and Glut1 at the lesion showed that the pattern of GFP^+^SC matched with that of Glut1+BV (Fig. 1C_3_). Co-labeling for Glut1 and GFAP showed that GFP^+^SC were localized in perivascular spaces between endothelial cells and astrocyte end-feet (Fig. 1E). SC within the BV lumen were never observed.

We used whole-mount immuno-labeling of clarified spinal cords to gain insight in SC-BV3D spatial organization. For these experiments, tdTomato^+^ SC were grafted remotely from the lesion as above, and their migratory behavior analyzed at 5 dpi. BV and SC were visualized by immuno-labeling against anti-IgG and anti-Tomato respectively. Light sheet imaging and 3D reconstruction revealed the dense spinal cord vascular network and confirmed that grafted SC, en route towards the lesion, were preferentially associated with BV in the spinal cord midline (Fig. 1F). TdTomato^+^ SC were either polarized along blood vessels (Fig. 1G,H,I) or extending processes from one vessel to another (Fig. 1J,K,L).

Whether SC moved along blood vessels was further confirmed by GFP^+^SC live-imaging 2 days post engraftment in the demyelinated spinal cords and host BV were labeled by intra-cardiac perfusion of rhodamine-lectin. Spinal cord whole mounts were maintained in appropriate physiological conditions, and areas containing GFP^+^SC were selected and video-recorded for 20 hours. Recordings confirmed that SC leaving the graft, reached for and became associated with BV, to migrate either in chains or as isolated cells, jumping from one BV to another one (MovieS1, Fig. S3A, blue empty arrowhead) or sliding along them (MovieS1, Fig. S3A, white arrowhead) as described above. While SC in chains glided along each other, individual cells escaped from the chain crawling along BV outer surface and merging further with other SC (Movie S2, Fig. S3B arrowhead).

To elucidate whether SC migrate *in vivo* in association with peri-vascular ECM, and/or whether they require direct interactions with endothelial cells, we performed immuno-electron microscopy (EM) on a novel series of GFP^+^SC grafted mice (Fig. 1M,N,O). EM analysis of DAB labeled GFP^+^SC confirmed their location in close contact with BV. Interestingly, the majority of the grafted SC was embedded within the peri-vascular ECM (Fig. 1M,N) between the vascular cells and astrocyte end-feet (Fig. 1O). However, unlike observed in the injured PNS(12), direct contact with vascular cells was never observed.

Temporal analysis of SC arrival at the lesion indicated that in lesions which had cells at early stages (3dpi), 62±9% of GFP^+^SC were closely associated with BV (Fig. 2A,C,G), while at later times (5dpi) only 32±5% of them were associated with BV (Fig. 2B,D,G). Immunohistochemistry for Glut1 and NF200 indicated that this transition correlated with a change in SC-BV to SC-Axon associations, with only 35±6% of GFP^+^SC associated with axons at 3dpi (Fig.2A,E,H), and 61±5% GFP^+^SC associated with axons at 5 days (Fig. 2B,F,H).

**Figure 2.**
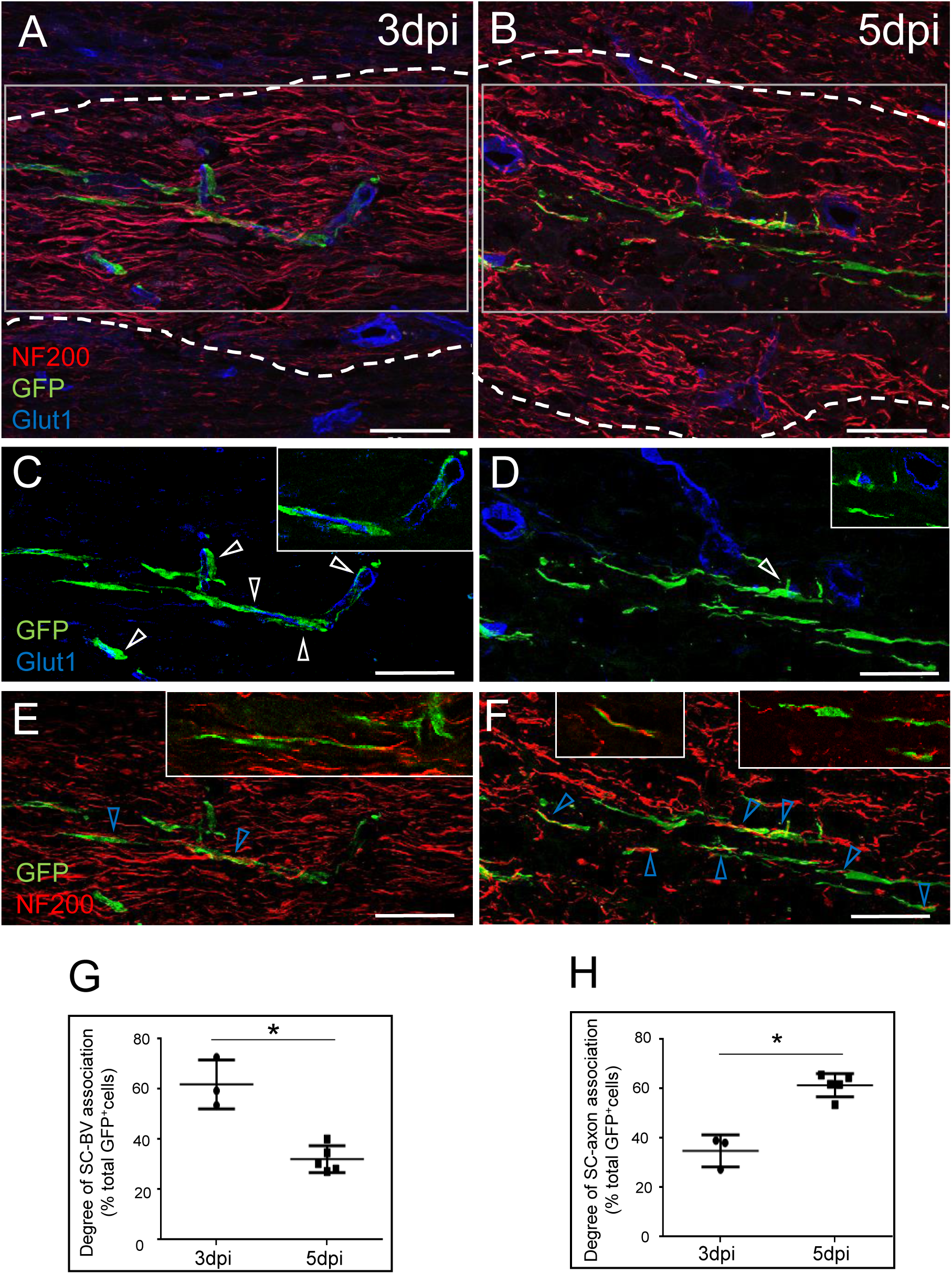
Schwann cells arrive at the lesion along BV and timely dissociate from them to contact demyelinated axons. (**A,C,E**) Upon arrival at the lesion at 3dpi, grafted GFP^+^SC are associated with BV. (**B,D,F**) In lesions at 5dpi, fewer SC are associated with BV but are aligned with NF200^+^axons. (**C-F**) Higher magnifications of GFP^+^ SC illustrating in (**C, E**) the temporal decrease of association of GFP^+^SC with Glut1-positive BV (white arrows) compared to (**E-F**) the progressive increase of GFP^+^SC association with NF200^+^ axons (blue arrow). **C, E** and **D, F** show separated colors of **A** and **B** respectively. Images represent confocal maximal projections of Z-stacks, while insets show only one confocal Z-plane. Scale bar 50 µm. (**G,H**) Quantification of SC associated or not, with BV at 3dpi (n=2) and 5dpi (n=8) (two-tailed Mann Whitney test p=0.035) (mean value ± SD of 3 independent experiments).

### LPC induced demyelination promotes angiogenesis and blood vessel remodeling

Since angiogenesis has been described in demyelinating diseases(36) and perivascular niche plays a role in remyelination(9), we examined the dynamics of BV in the absence of SC grafting. We found an increase in BV density in lesions of non-grafted spinal cords (Fig. 3B-E), compared to PBS-injected spinal cords (Fig. 3A). Glut1-positivity increased from day 3dpi up to 7dpi with a peak at 5dpi compared to PBS injected controls (Glut1^+^area in Lesion/PBS (n=15); 1dpi (n=4): 1±0.05; 3dpi (n=4): 1.8±0.5; 5dpi (n=4): 2.2±0.4; and 7dpi (n=7): 1.6±0.5; values are expressed as the ratio to the PBS mean) (Fig. 3I). This transient increase in BV density in LPC lesions was partially due to increased proliferation of endothelial cells based on the number of Glut1+cells, expressing Ki67 (Ki67^+^Glut1^+^ cells/mm2: 1dpi (n=4): 12.7±7 (p=0.004), 3dpi (n=4): 7.8±7 (p=0.001), 5dpi (n=4): 6.5±6 (p=0.002), 7dpi (n=7): 2.9±4 (p=0.001))(Fig. 3A-F,J, Fig. S4, S5, Supplementary note 1), pointing to an increase of newly formed vessels as a specific reaction to demyelination in addition to BV remodeling. Immuno-labeling for FN and collagen 4 showed that LPC-induced angiogenesis/BV remodeling was concomitant with changes in perivascular ECM (Fig. S6).

**Figure 3.**
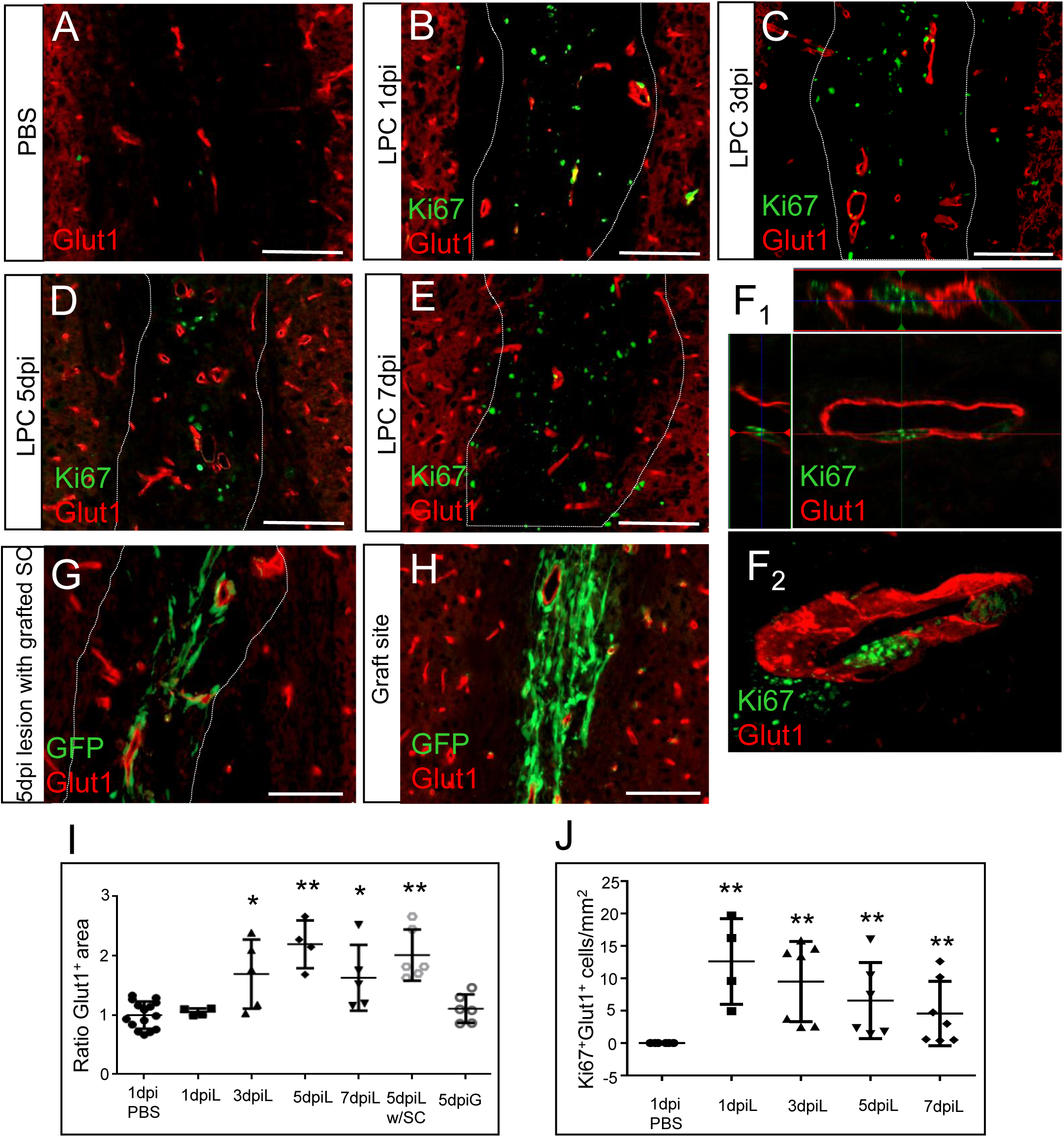
Dynamics of the vascular network in response to LPC demyelination. (**A-F**) Immunohistochemistry for Glut1^+^ (red) at 1dpi (**B**), 3dpi (**C**), 5dpi (**D**) and 7dpi (**E**) illustrates increased vascularization in LPC lesions compared to control (**A**). (**F_1_**) Orthogonal and **(F_2_)** 3D views show proliferating Glu1^+^Ki67^+^ endothelial cells. (**G,H**) Immunohistochemistry for Glut1^+^ at lesion site (**G**) and graft site (**H**) in SC grafted mice at 5dpi. Dashed lines highlight the lesion border. (**I**) Evaluation of Glut1^+^area shows a significant expansion of BV from 3dpi-7dpi, with a peak at 5dpi, which is similar to that observed in lesions of grafted animals, and no increase in Glut1^+^ area at the graft site. One-way ANOVA p<0.001, F(4,28)=10.96; two-tailed Mann Whitney test with PBS group: 3dpi (p=0.019); 5dpi (p=0.0005); and 7dpi (p=0.01). (**J**) The number of Ki67^+^/Glut1^+^cells increases at 1dpi preceding that Glut1^+^area, and declines thereafter (one-way ANOVA p=0.007, F(4,25)=4.48; two-tailed Mann Whitney test with PBS group: 1dpi (p=0.004), 3dpi (p=0.001), 5dpi (p=0.002), 7dpi (p=0.001). Data are expressed as mean value ± SD of control (PBS) of 3 independent experiments, PBS group (n=15) 1dpi (n=4), 3dpi (n=4), 5dpi (n=4), 7dpi (n=7), 5dpi without SC (n=6), 5dpi in SC graft site/PBS (n=6). **means p<0.01 and *means p<0.05. Scale bar 100 µm.

We examined whether the presence of exogenous SC could influence the formation of new blood vessels. We found no difference in Glut1^+^area in lesions of SC-grafted mice (5dpi: 2.2±0.4) compared to lesions of non-grafted ones at 5dpi (Glut1^+^area lesion of non-SC grafted mice/PBS at 5dpi in: 2±0.4)(Fig. 3G,I). Moreover, no change in Glut1^+^area was observed at the SC graft site compared to PBS injected mice at 5dpi (5dpi in SC graft site/PBS: 1.1±0.2)(Fig. 3H,I).

### Myelin-associated EphrinB3 as candidate to guide Schwann cells in the CNS

Since no effect of exogenous SC on vascularization was observed, we hypothesized that SC guidance within the CNS by SC-BV association and myelin repulsion could be due to some receptor-ligand mediated mechanism. We previously demonstrated that SC are repulsed by CNS myelin, partially mediated by the axonal repellent MAG, suggesting that other myelin components might be involved in this repulsion(20).

Thus, we speculated that other axonal growth inhibitors were able to induce the same effect in SC. Of interest, EphrinB3 is expressed by CNS myelin and inhibits axonal growth and oligodendrocyte differentiation(25, 26), and SC express several Eph receptors (28). Interestingly, immature stages of the SC lineage such as boundary cap cells and SC precursors, have a greater capacity to migrate through white matter compared to more mature SC stages(37, 38) present lower expression of another EphrinB3 receptor, the EphB6(39). In support of these data, we performed gene profiling by RNA sequencing, comparing aged-matched wild-type and developmentally arrested SC lacking Krox20 (Krox20^null^ SC)(40)(considered at a more immature-like stage) revealed a dysregulation in the transcript levels of a variety of molecules among which Eph receptors were particularly expressed at lower level. While EphA4, EphB1, EphB2, EphB3, EphB4 and EphB6 transcripts were expressed in wild-type SC, EphA4, EphB1 and EphB6 were significantly down-regulated in Krox20^null^ SC (Fig. S7). Hence, we hypothesized that EphA4, EphB6, and EphB1 could be involved in SC poor mingling with CNS white matter, and favorable association with BV.

### The EphrinB3 present in myelin is able to bind and activate EphrinB3-receptors in Schwann cells

We first confirmed that SC have the molecular machinery to bind myelin-associated EphrinB3. Immunohistochemistry showed that purified mouse SC can bind EphrinB3 *in vitr*o, and that these cells express the three EphrinB3 receptors: EphB6 (Fig. 4A), EphA4 (Fig. 4C) and EphB1 (Fig. 4D). The presence of each receptor was corroborated by Western blot (Fig. 4B). Incubation of SC with pre-clustered EphrinB3 by fluorescent anti-Ig and orthogonal views revealed that the three receptors were able to bind clustered EphrinB3 on SC surface leading to the formation of large signaling clusters as previously described(41) (Fig. 4A,C,D).

**Figure 4.**
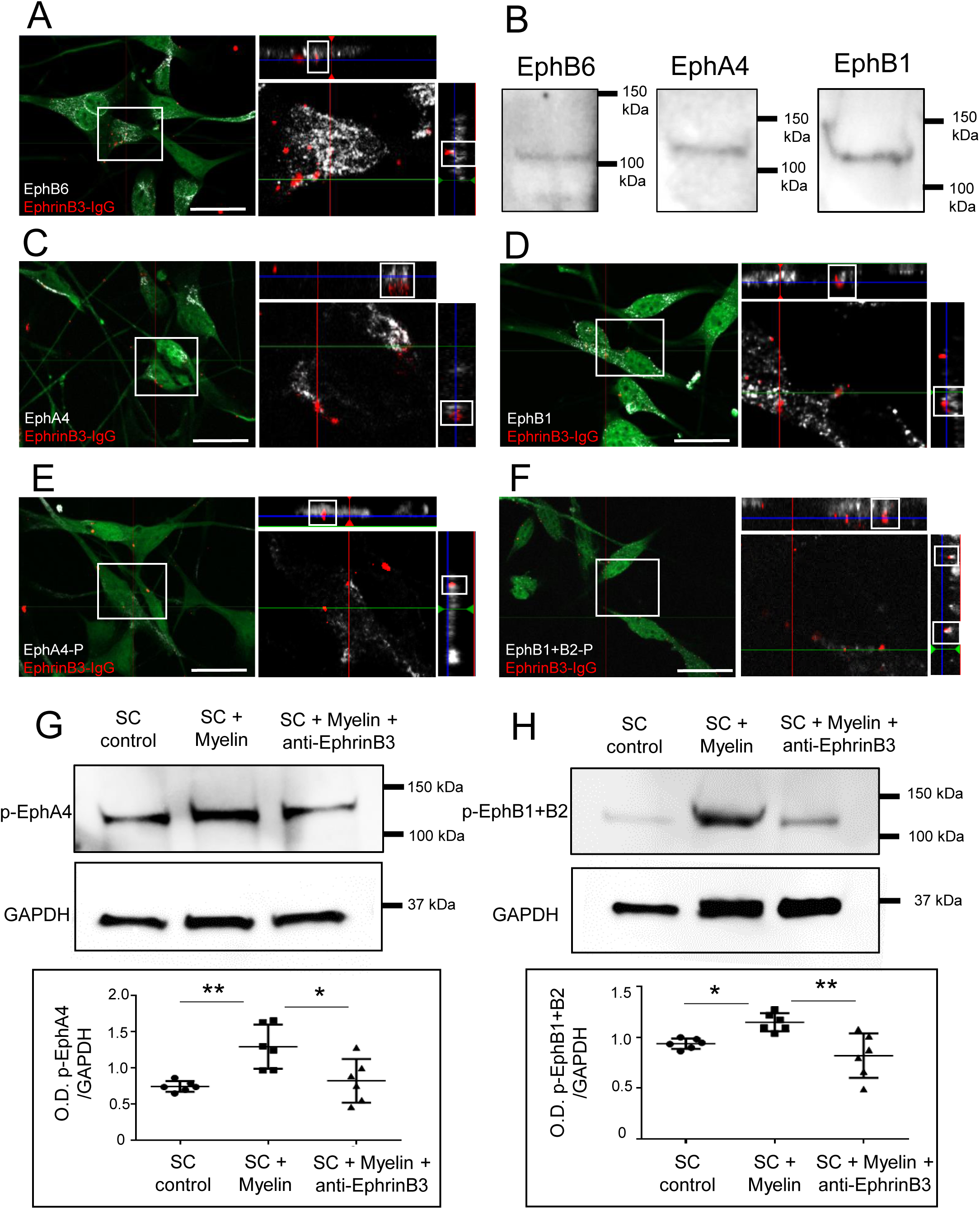
EphrinB3 binds and activates EphA4, EphB1 and EphB6 receptors in SC. (**A, C, D**) Orthogonal views of SC activation with clustered-EphrinB3 shows that EphrinB3 binds EphB6 (**A**), EphA4 (**C**) and EphB1 (**D**) receptors on SC. (**B**) Western blot confirming the presence of these receptors in SC. (**E,F**) Bound EphrinB3 activation of EphA4 and EphB receptors viewed by immunodetection of phosphorylated forms. (**G,H**) Western blot illustrating that SC incubation with myelin increased the phosphorylation of EphA4 (**G**) and EphB1+B2 (**H**) which was not the case when myelin was previously blocked by anti-EphrinB3 (O.D. p-EphB1+2/GAPDH: one-way ANOVA p=0.003, F(2,15)=8.57, followed by a Tukey’s multiple comparison test); and O.D. of p-EphA4/GAPDH: one-way ANOVA p=0.0036, F(2,15)=8.44, followed by a Tukey’s multiple comparison test). Data are expressed as ratio of the optical density (O.D.) of the bands (mean values ± SD) from 3 independent experiments, control (n=6), myelin (n=6), myelin+anti-EphrinB3 (n=6). Scale bar 20 µm.

Eph receptors are activated by auto-phosphorylation of specific tyrosine residues(42). To verify the ability of EphrinB3 to induce forward signaling, activation of Eph receptors on SC was assessed. Immunocytochemistry illustrates binding of EphA4 with clustered EphrinB3, and its activation by the ligand (Fig. 4E). EphB6 is a recognized Kinase-defective receptor, which can form a hetero-receptor complex with EphB1 receptor and undergoes trans-phosphorylation(43). Immunohistochemistry validated the presence of phospho-Y594 in EphB1 (and EphB2) receptor, and its interaction with bound EphrinB3 in SC (Fig. 4F).

To test the hypothesis that myelin associated EphrinB3 is able to induce the activation of EphB and EphA4 receptors, we incubated purified SC during 30min with myelin protein extracts or myelin protein extract previously incubated with anti-EphrinB3 known to block specifically its activity(26). SC were collected and blotted with an antibody against the phosphorylated EphA4 (Fig. 4G), and an antibody against phosphorylated EphB1+B2 (Fig. 4H). The presence of myelin increased significantly the phosphorylated forms of both receptors compared to the housekeeping GAPDH, but this signal increase was not observed when the myelin extracts were pre-incubated with anti-EphrinB3 antibody before activation ((O.D. of p-EphB1+2/GAPDH control: 0.93±0.05, myelin: 1.1±0.08, myelin+anti-EphrinB3: 0.82±0.21) and O.D. of p-EphA4/GAPDH control: 0.74±0.07, myelin: 1.29±0.30, myelin+anti-EphrinB3: 0.82±0.30). Blocking the epitopes of EphrinB3 in myelin neutralized the Eph receptor activation effect of myelin on SC (Fig.4G,H).

Hence, these observations evidenced the ability of SC to bind and respond to the presence of EphrinB3 *in vitro*.

### CNS myelin inhibits Schwann cell adhesion and spreading, and this effect is partly mediated by EphrinB3 through EphA4 and EphB6 receptors

Next, we examined whether EphrinB3 could contribute to SC-myelin repulsion. We performed a blocking receptor assay, pre-incubating purified GFP^+^SC with unclustered EphrinB3-Fc molecules before seeding them for 3 hours onto myelin protein extract or PBS (Fig.5A), and evaluated both SC adherence and polarization (Fig.5B). Control Fc-preincubated SC showed less adhesion to myelin, measured as the ratio of adhered SC to myelin compared to PBS coated surfaces. SC also extended fewer processes, based on a higher ratio of round cells over the total adhered SC on myelin compared to control. Blocking EphrinB3 receptors on SC by soluble EphrinB3-Fc hindered myelin repulsion, resulting in a significant improvement of SC adhesion to myelin-coated surfaces, and ability to extend more processes (fewer round cells) on myelin than those pre-incubated only with soluble Fc (Ratio of adhered cells Myelin/PBS; Fc-preincubated SC on myelin: 0.4±0.2; EphrinB3-preincubated SC on myelin: 0.8±0.2. Ratio of round cells Myelin/PBS: Fc: 2.9±1; EphrinB3: 1.5±0.5). (Fig.5A,B).

To determine if the reduction in the number of SC adhering to myelin was due to cell death, we assessed cell viability comparing the percentage of caspase3^+^ nuclei or Hoechst^+^pycnotic nucleiin SC incubated with PBS or myelin for 3h. SC exhibited a typical bipolar morphology and no difference in the percentage of caspase3^+^nuclei or Hoechst pycnotic nuclei between control and myelin-treated groups were observed (PBS: 4.4±1.6‰, Myelin: 4.0±1.7‰, n=6). These data corroborate our previous results demonstrating that short incubation time with myelin does not induce apoptosis(20).

SC adhesion to surfaces coated with EphrinB3-Fc was reduced compared to Fc as control in a dose-dependent manner (Ratio of adhered cells EphrinB3/Fc; 2.5µg/mL: 0.9±0.3, 10µg/mL: 0.7±0.07 and 20µg/mL: 0.5±0.2. Ratio of round cells EphrinB3/Fc: 2.5µg/mL: 1.2±0.4, 10µg/mL: 1.8±0.4 and 20ug/mL: 3±0.9)(Fig. 5C). Moreover, the increased number of round SC indicated that SC were less polarized on EphrinB3 coated surfaces compared to control (significance reached with high concentrations of EphrinB3) (Fig. 5C).

**Figure 5.**
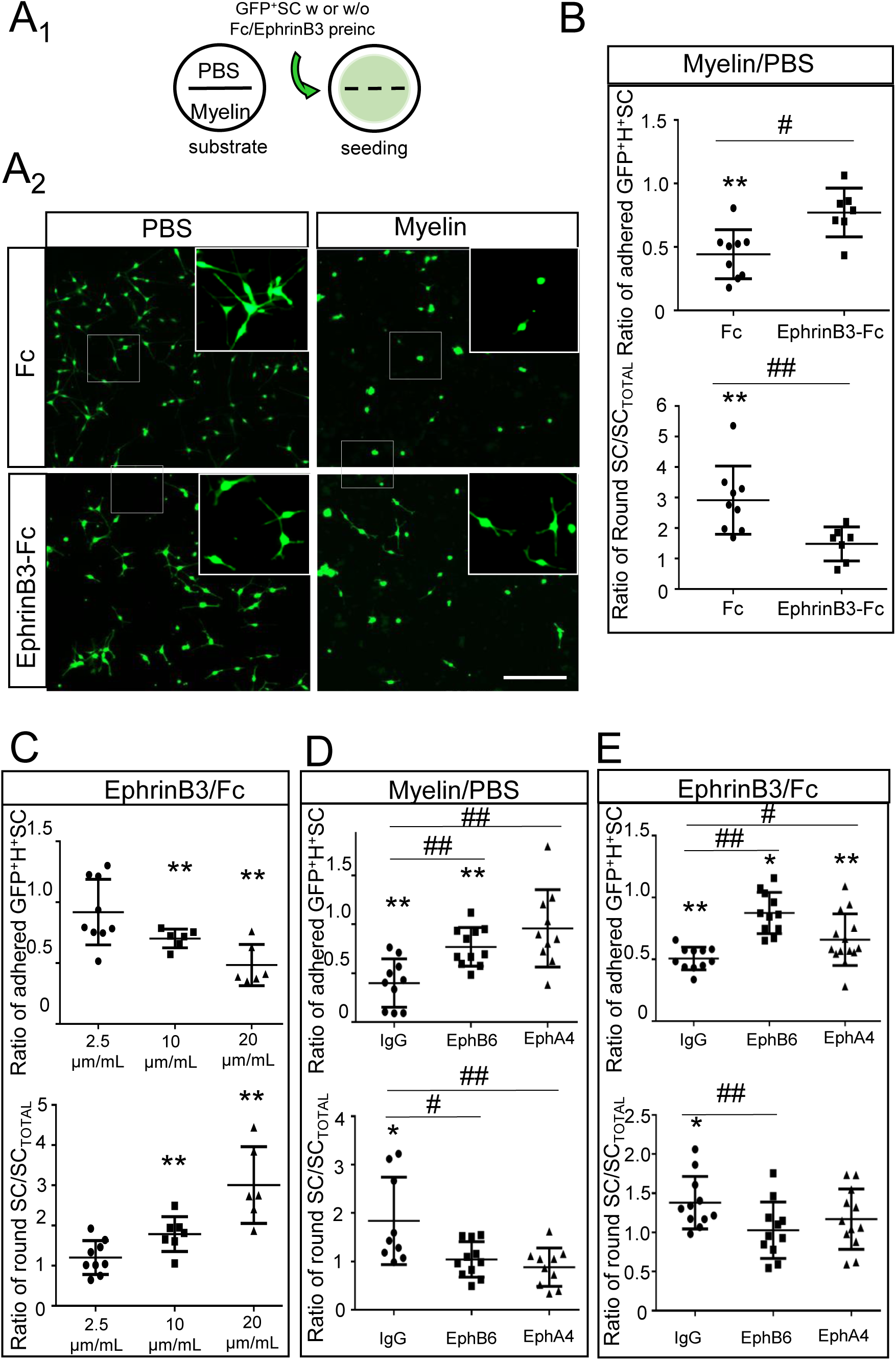
EphrinB3 mediates myelin-inhibition on SC adhesion and polarization through EphA4 and EphB6 receptors. (**A_1_**) Diagram of the adhesion assay. Coverslips divided by silicon strips were coated with PBS or myelin in each half. After strip removal, SC pre-incubated with Fc or EphrinB3 were seeded homogenously on the coverslip. (**A_2_**) Unclustered EphrinB3 partially reverted myelin-induced inhibition of SC adhesion and spreading, scale bar 200 µm. (**B**) Quantification of the ratio of adhered or round GFP^+^SC on Myelin over PBS after pre-incubation with unclustered EphrinB3 or Fc. (**C**) SC seeded on substrate coated with increased concentrations of EphrinB3, adhered and spread less in a dose dependent manner respect to the intra-coverslip control substrate (Fc). EphrinB3/Fc: 2.5µg/mL, 10µg/mL, and 20µg/mL. (**D, E**) Quantification of the number of adhered and polarized SC pre-incubated with anti-EphB6 or anti-EphA4 (extracellular domain). Pre-incubation with anti-EpB6 and anti-EphA4 improved the number of adhered and polarized SC on Myelin (**D**) and EphrinB3 (**E**) compared to PBS and Fc respectively. Pre-incubation with IgG as control showed similar results than non-pre-incubated cells (**B, C**). Data are expressed as ratios (mean values ± SD form 3 independent experiments) of Myelin or EphrinB3 surfaces compared to intra-coverslips control non-coated surfaces (PBS) or with equimolar concentration of Fc respectively. In **B, C, D, E** */** are used for comparison of a group with its hypothetical mean: 1 by one sample two-tailed t-test, and #/## for comparison between two different groups by two-tailed Mann Whitney test (n=7-11 per group) */# means p<0.05; **/## means p<0.01.

EphrinB3 behaves as a dependence receptor, which can trigger cell apoptosis(44). To analyze the potential effect of EphrinB3 on Schwann cell survival, we assayed cell viability as above. Incubation of adhered SC with clustered EphrinB3, or Fc as controls, for 3h and 24h did not induce any morphological difference, and the number of H^+^pycnotic or caspase3^+^nuclei between Ephrin-treated and control SC remained equivalent (3h treatment (n=6): Fc: 4.4±2‰, EphrinB3: 4.4±2‰; 24h post-treatment (n=8): Fc: 9.7±4‰, EphrinB3: 10.4±3‰). Adhesion assays were always performed in serum-free medium, compromising SC proliferation. These results imply that the reduction in the number of attached and spread SC to the different substrates results from their preferential adhesion rather than a toxic effect of the treatment.

To validate the involvement of EphA4 and EphB6 receptors in SC-EphrinB3 response, we performed the myelin and EphrinB3 adhesion assay in the presence of antibodies that specifically interfere with EphA4 or/and EphB6 receptors, to block SC response. Blocking EphA4 or EphB6 receptor with excess anti-EphA4 or anti-EphB6 respectively improved SC adhesion and spreading to myelin compared to PBS (Fig. 5D) or to EphrinB3 compared to Fc (Fig. 5E). Ratio of adhered cells Myelin/PBS: IgG: 0.4±0.2; anti-EphB6: 0.9±0.4, anti-EphA4: 0.8±0.2. Ratio of adhered cells EphrinB3/Fc: IgG: 0.5±0.09, anti-EphB6: 0.9±0.2, anti-EphA4: 0.6±0.2. Ratio of round cells Myelin/PBS: IgG: 1.8±0.9, anti-EphB6: 1.0±0.4, anti-EphA4: 0.8±0.4. Ratio of round cells EphirnB3/Fc: IgG: 1.4±0.3, anti-EphB6: 1.0±0.4, anti-EphA4: 1.2±0.4, data are expressed as ratio (mean values ± SD) of Myelin or EphrinB3 surfaces compared to intra-coverslips control non-coated surfaces (PBS) or with equimolar concentration of Fc respectively. However, no synergistic/additive effect on adhesion was observed when antibodies to both receptors were combined (Myelin/PBS: anti-EphA4 + anti-EphB6: 0.8±0.2(n=11), *p*(IgG vs anti-EphA4 + anti-EphB6)=0.003; EphrinB3/Fc:anti-EphA4 + anti-EphB6: 0.8±0.3(n=10), *p*(IgG vs anti-EphA4 + anti-EphB6)=0.008) two-tailed t-student test).

### EphrinB3 increases Schwann cell adhesion and migration on the perivascular ECM component, fibronectin, via integrin ß1

Eph/ephrin can modulate cellular pathways by regulating cell adhesion, either positively or negatively, depending upon the cellular context(45). Thus, positive regulation of SC adhesion to ECM and FN might have consequences on their migration capacity on this substrate(32, 33). As our *in vivo* study indicated that the grafted SC were embedded in perivascular ECM, and angiogenesis is a response to demyelination, we studied the effect of EphrinB3 on SC-ECM binding, in particular FN, that favors SC migration(10). Unlike SC repulsion by EphrinB3-coated surfaces, EphrinB3 improved SC adherence and process expansion on FN coated surfaces compared to those coated with FN and Fc (Fig. 6A). Ratio of adhered cells: EphrinB3 (10µg/mL): 0.7±0.07, EphrinB3 (10µg/mL) + FN (2µg/cm^2^): 1.07±0.1. Ratio of round cells: EphrinB3 (10µg/mL): 1.8±0.4, EphrinB3 (10µg/mL) + FN (2µg/cm^2^): 0.82±0.3 (data are expressed as ratio (mean values ± SD) respect to Fc-non FN coated surfaces).

**Figure 6.**
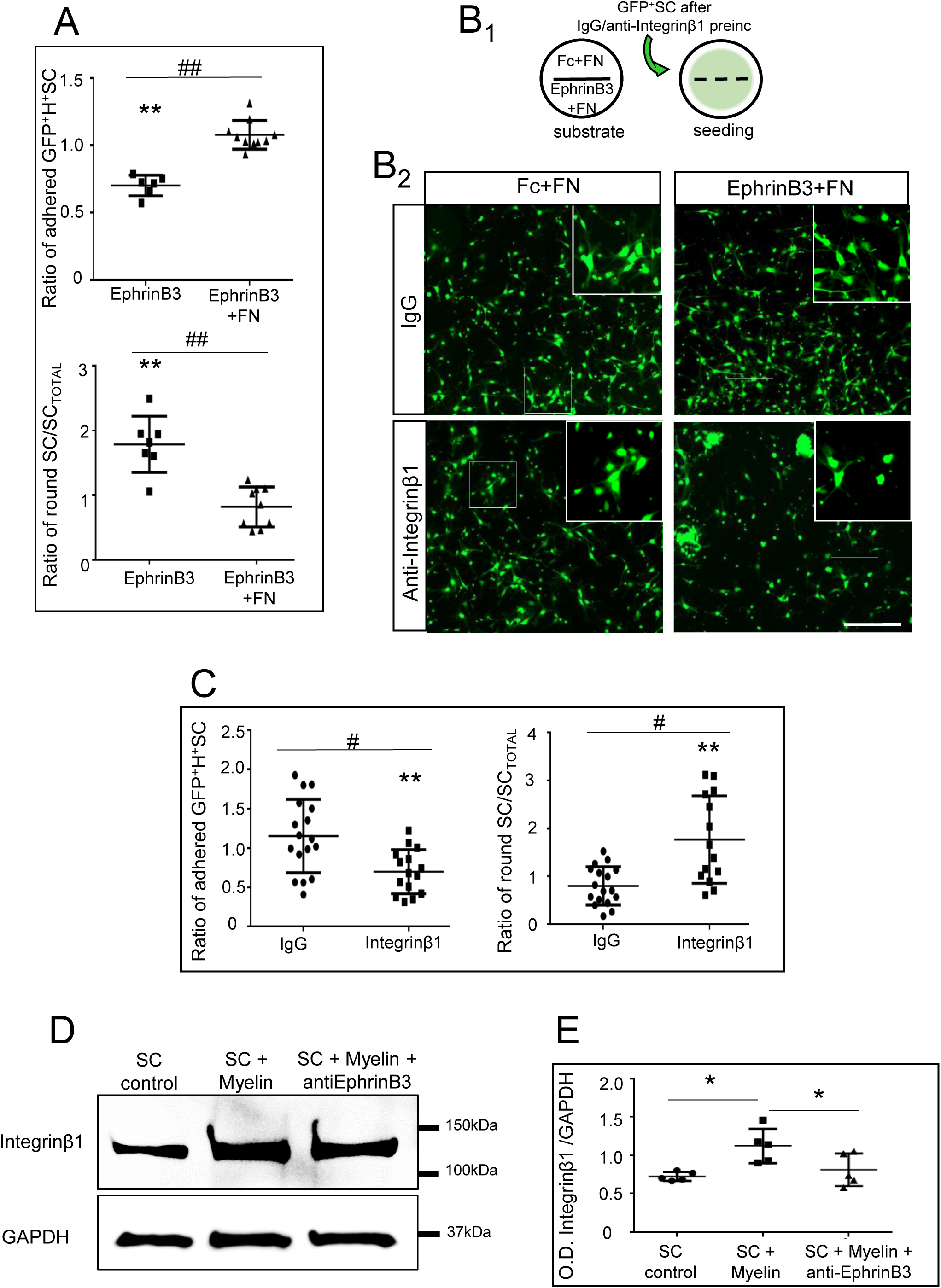
EphrinB3 improves SC adhesion and spreading on FN coated surfaces. (**A**) Quantification of the number of adhered and polarized SC seeded on substrate coated with EphrinB3 with or without FN. Data are expressed as ratio respect to Fc-coated surfaces. EphrinB3 (10µg/mL), EphrinB3 (10µg/mL)+FN (2µg/cm^2^), n=7 per group. (**B_1_**) Diagram of the adhesion assay on FN. Coverslips divided by silicon strips were coated with Fc+FN (control) or EphrinB3+FN in each half. After strip removal, cells pre-incubated with IgG or anti-Integrinβ1 were seeded homogenously on the coverslip. (**B_2_**) Blocking Integrinβ1 decreased SC adhesion and polarization on EphrinB3+FN compared with Fc+FN, scale, bar 200µm. (**C**) Quantification of SC adhesion and polarization on EphrinB3 when SC are pre-incubated with IgG or anti-integrinß1, n=15-17 per group. (**D-E**) Western blot analysis shows increased expression of Integrinβ1 when SC are activated by myelin extracts. One-way ANOVA p=0.011, F(2,12)=6.62, followed by Tukey’s multiple comparisons test p<0,05, n=5 per group. In **A, C**\*/** are used for comparison of a group with its hypothetical mean: 1 by one sample two-tailed t-test, and #/## for comparison between two different groups by two-tailed Mann Whitney test.*/# means p<0.05; **/## means p<0.001.

The main FN-binding receptor on SC is the integrin heterodimer α5β1(46), and the integrin family is involved in the Eph/ephrin response(31, 32). To question whether integrinß1 could mediate the mechanism by which EphrinB3 regulates SC-FN binding, we performed an integrinβ1 interfering assay prior to SC adhesion on FN+EphrinB3 (Fig. 6B,C; IgG: 1.1±0.4; anti-integrinβ1: 0.7±0.3; data are expressed as ratio over control (Fc+FN substrate)). Anti-integrinβ1 pre-incubation with a specific blocking antibody(47) consistently prevented the increased SC adhesion to FN+EphrinB3 coated surfaces compared to the FN+Fc coated ones. Moreover, Western blot showed that incubation of SC during 30 min with myelin protein extracts (100µg/mL) increased expression of Integrinβ1 and this increase was prevented by pre-incubation of the myelin protein extract with anti-EphrinB3 (O.D. of Integrin β1/GAPDH in control: 0.72±0.05, myelin: 1.12±0.22, and myelin+anti-EphrinB3: 0.81±0.21)(Fig.6D, E). This suggests that myelin associated EphrinB3 induced integrinβ1 expression, which is involved in the increased ECM adhesion induced by EphrinB3.

We questioned whether this induced SC adhesion to FN could have some implication in their ability to migrate on FN by using the agarose drop assay(48). SC were seeded on FN in the presence of EphrinB3 or Fc, and their migration when exiting the agarose drop, was followed by time-lapse video-microscopy. Significantly more SC migrated out of the drop when sections were coated with FN+EphrinB3 (Fig.7B,C) compared to control (Fig. 7A,C). Moreover, based on their maximal distance of migration, SC migrated significantly further on FN+EphrinB3 compared to control (Fig. 7D).

**Figure 7.**
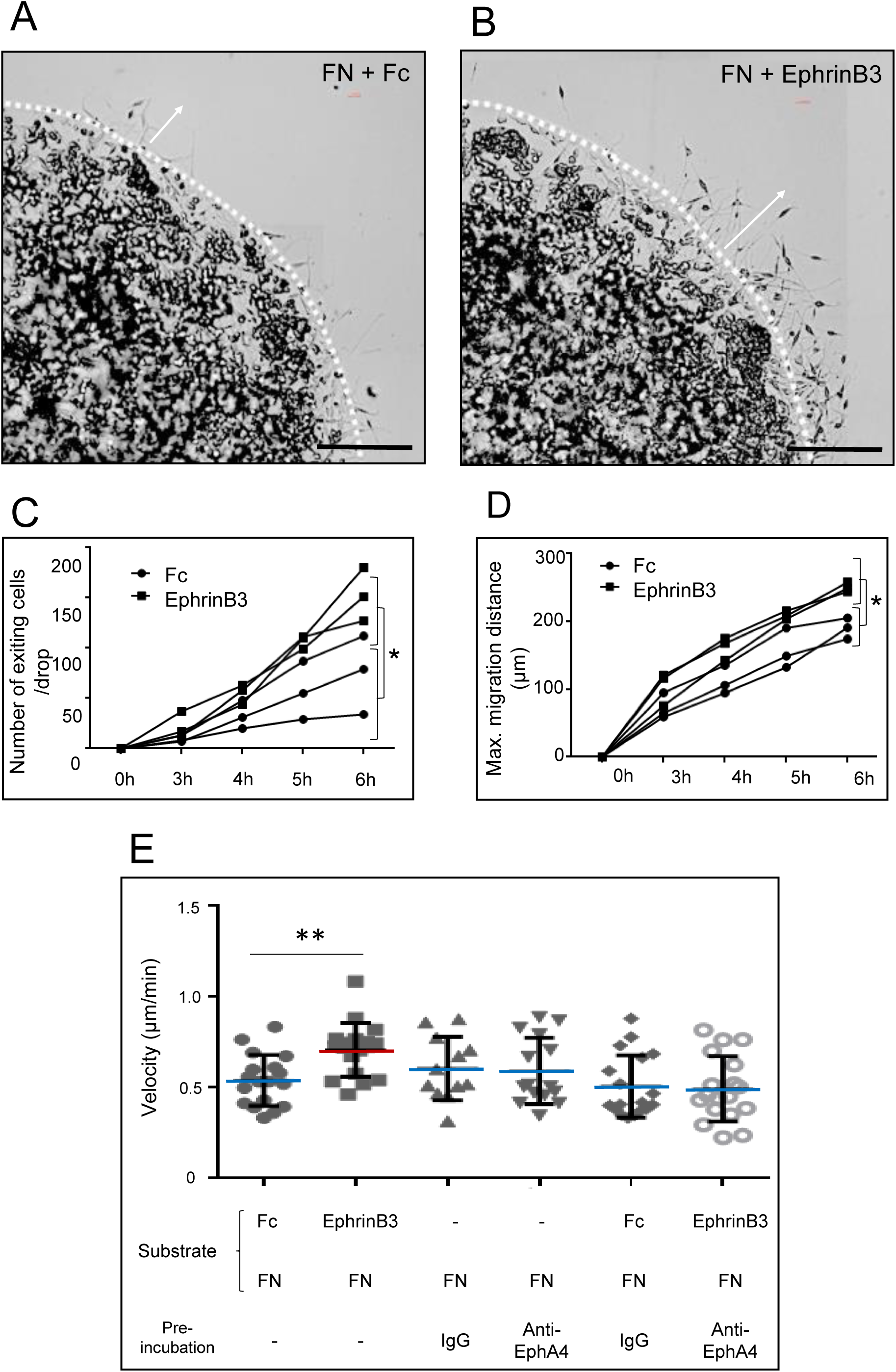
EphrinB3 improves SC migration on FN coated surfaces. (**A,B**) Exit of SC entrapped in an agarose drop and seeded on FN+Fc (**A**) and FN+EphrinB3 (**B**), scale bar 200µm. (**C**) More SC exit the agarose drop 5h post-seeding (two-way ANOVA with repeated measures: p=0.03, F(1,4)=10.24). (**D**) SC migrate over longer distances from the drop-edge 4h post-seeding (two-way ANOVA with repeated measures: p=0.02, F(1,4)=12.74) on FN+EphrinB3 coated surfaces (n=3 per group). Graphs represent the values of separate experiments (mean±SD). (**E**) Velocity of single SC was measured in different conditions. Only SC seeded on FN+EphrinB3 increase significantly their migration speed. SC velocity (μm/min) in conditions of Fc+FN coating (n=18), FN+EphrinB3 coating (n=18), pre-incubation with IgG on FN coating (n=12), pre-incubation with anti-EphA4 on FN coating (n=15), pre-incubation with IgG on FN+Fc coating (n=18), pre-incubation with anti-EphA4 on FN+EphrinB3 coating (n=18). This increase is reverted when SC are pre-incubated with anti-EphA4 (One-way ANOVA p=0.002, F(5,93)=4.1; two-tailed Mann Whitney test p=0.002 between control FN+Fc and FN+EphrinB3 group). In **E** (n) represents tracked single cells of 3 different experiments repeated independently. Data are expressed as mean values ± SD. Single cells were tracked from three different independent experiments, ** means p<0.01

Finally, we performed interference experiments to analyze the migratory behavior of SC incubated or not with anti-EphA4 on FN substrate alone or FN+Fc or FN+EphrinB3. Analysis of the migration velocity of single cells traced by video microscopy on FN confirmed that SC migrate significantly faster on FN+EphrinB3 substrate, compared to Fc+FN control (Fig. 7E). This increased speed was counteracted when SC were pre-incubated with anti-EphA4 compared to control IgG. Moreover, data confirmed that pre-incubation with IgG or anti-EphA4 did not impair SC speed or pattern of migration on FN whether seeded over Fc or EphrinB3 (Fig. 7E).

### *P*re-treatment with anti-Eph4 promotes Schwann cell to mingle more with myelin *in vivo*, and reduces their interaction with blood vessels

*In vitro* experiments established that EphrinB3 had a dual effect, impairing SC interaction with myelin but improving their interaction with FN via increased integrinß1 expression. Moreover perturbation experiments indicated that anti-EphA4 treatments did not affect SC migratory behavior. To examine whether EphrinB3 plays a role in their integration/migration into CNS white matter and/or their interaction with BV *in vivo*, we interfered with EphrinB3 by blocking EphA4, and examined SC interactions with BV and myelin. Since no synergic effect was observed *in vitro* when blocking two different receptors, EphA4 appeared to be the candidate of choice. Pre-incubation of SC with the functional anti-EphA4 blocking antibody prior transplantation as above, disrupted SC interaction with BV (Fig. 8A-C compared to D-F), evaluated by the number of SC-BV association (Fig. 8O), and improved their mingling with myelin along their pathway of migration (Fig. 8G,H vs 8I,J). While 67% of SC was associated with BV in the control group, only 45% did so in the anti-EphA4 treated group (Fig.8O). Moreover, for the same graft-lesion distance, lesion size and amount of grafted cells, more GFP^+^SC were found in the lesion site at 5dpi in animals grafted with IgG treated-SC (Fig. 8C,K,L) compared to those grafted with anti-EphA4 treated SC (Fig. 8E,M,N,P). Thus, anti-EphA4 treatment reduced the capacity of SC to progress efficiently along vessels and/or enhanced their sensitivity to other myelin inhibitors.

**Figure 8.**
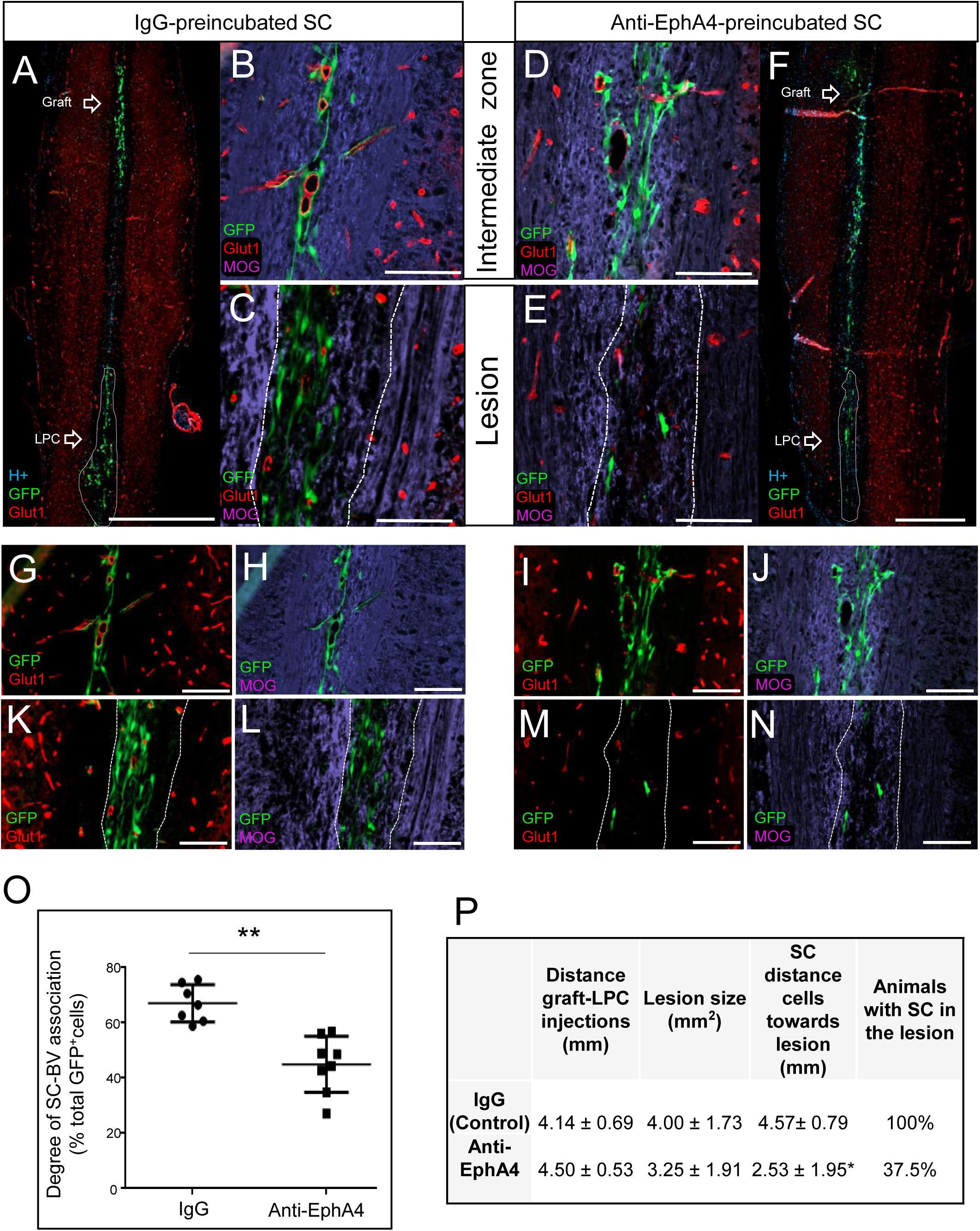
Pre-incubation of SC with anti-EphA4 reduces their migration along BV after transplantation in the demyelinated spinal cord. (**A**) GFP^+^SC pre-incubated with IgG migrate from the graft along the midline in association with Glut1^+^BV (**B,G**) but avoid MOG^+^ myelin (**B,H**) before arrival at the lesion at 5dpi (**C,K,L**). (**F**) Anti-EphA4-preincubated GFP^+^SC exiting the graft mingle more with myelin (**D, J**), associate less with BV (**D,I**) and frequently fail to reach the lesion site at 5dpi (**E,M,N**). (**O**) Quantification of SC in association or not, with BV show significant differences between control (IgG) (n=7) and anti-EphA4 pre-incubated SC (n=8) with fewer cells associated with BV after anti-EphA4 pre-incubation (Mann Whitney test, p=0.0003). (**P**) Quantification of the extent of migration of grafted GFP^+^SC. The reduced extent of migration by SC pre-incubated with anti-EphA4 prior grafting, compared to control SC, correlates with a reduced percentage of animals in which grafted SC are recruited by the lesion. Dashed lines delineate lesions. Data are expressed as mean values ± SD; * means p<0.05. Scale bar 100 µm, except for A and F: 1000 µm.

## DISCUSSION

Compelling evidences indicate that SC, whether recruited from the periphery or generated from resident progenitor cells, invade the CNS to remyelinate CNS axons after demyelination(5-9), having an obvious impact on clinical recovery(1, 4). However, in spite of their presence within the CNS, SC intrusion of the CNS is restricted to BV proximal to the PNS, and the mechanisms regulating PNS-CNS transgression after myelin injury, remain elusive. To gain insights into PNS-CNS poor interface, we studied SC behavior when confronted with CNS myelin and BV *ex vivo* and *in vivo* in demyelinating conditions. Using classic, live, 3D imaging as well as electron microscopy, we provide solid evidences that SC use the vascular scaffold to migrate within the adult demyelinated CNS. This phenomenon is doubly modulated by increased angiogenesis and perivascular ECM in response to demyelination, as well as by SC restricted migratory capacity by CNS myelin. In particular, the CNS myelin specific component, EphrinB3, negatively regulates SC adhesion to-and spreading on myelin, while enhancing SC adhesion to perivascular ECM and promoting their migration along them.

Extensive intercellular communication and coordinated interaction between the vascular and nervous systems(49, 50) results in a functional neurovascular unit that contributes to wound healing(12), immune response(51) and embryonic development(52). Recently, a new role in supporting long distance migration of different kinds of cells within the nervous systems was attributed to BV, both in the developing and adult nervous system(13, 14) as well as under pathological conditions(12, 15). So far, despite the increasing number of cells guided by BV, the role of these structures in guiding SC within the CNS to participate in CNS repair was not explored. While SC are known to promote endothelial cell migration and angiogenesis(53) as well as to use BV as scaffold during peripheral nerve regeneration(12), our work demonstrates that this SC-BV interface extends to the CNS and is of relevance to their contribution to CNS repair.

We show that BV are a favorable substrate for exogenous SC and guide them towards demyelinating lesions. BV serve as scaffold for SC as soon as they leave the graft and along their path until arrival at the lesion. Along this path, SC are organized in chains going from one BV to another. At their arrival in the lesion, SC embedded in perivascular ECM become more randomly dispersed and this dispersal faithfully overlaps with BV expansion with time (Fig. 1). This suggests that angiogenesis, a common response to demyelination and inflammation, via expansion of the vascular network and ECM increase, facilitates SC entry sites and dispersal in the lesion. Since SC remyelination has shown to have an impact in the nerve function recovery(4), the clinical impact of angiogenesis in remyelination should not be overlooked.

While associated with BV at their arrival at the lesion at 3dpi, SC detached from BV to contact and align with the demyelinated axons at 5dpi. This change in substrate association may result from signals arising from the axons that trigger their differentiation into more mature SC as a first step to myelin repair. The absence of SC away from their narrow path of migration between the graft and lesion as well as their progressive increase in number in the lesion, point to their specific recruitment by the lesion most likely involving attractant signals yet to be defined. Although most of our observations were performed with exogenous SC, induction of demyelination in Krox20 cre Rosa YFP mice, hints that PNS-derived SC(7) triggered to enter the spinal cord by demyelination, use BV as scaffold to reach the lesion during spontaneous repair. SC are generated also from CNS progenitors in response to demyelination(8,9) opening the possibility that these CNS progenitor-derived SC may also use BV to migrate towards more distant injured sites.

In physiological conditions, the vasculature network, consisting of endothelial cells and pericytes surrounded by ECM, is rather dynamic, and angiogenesis is activated in response to damage(54). As part of this structure, ECM undergoes constant remodeling(55), as also observed in MS and animal models, even before symptoms appear(56). We show that injection of the lipid-toxic agent, LPC, in the absence of grafted SC, induced proliferation of endothelial cells in addition to expansion of the vasculature area. As BV remodeling is known to correlate with a change in ECM, we analyzed this impact in our paradigm. We found a prominent increase in FN and collagen 4 expressions, two main ECM components, which provide excellent substrates for SC migration. FN re-expression also affects directly OPC migration and differentiation and therefore, was proposed as a scaffold required to complete remyelination(57). Likewise, FN promotes SC mobility(10) indicating that the observed changes in ECM composition could stimulate SC migration along BV.

BV also support SC migration in the injured PNS. However, in those circumstances, SC create direct contacts via protrusion with endothelial cells to migrate from one nerve stump to the other or *in vitro* when grown in 3D, suggesting that these protrusions constitute mechanical means to propel SC migration along BV in a tight environment(12). Although we found SC associated with BV along their migrating route, direct contact with endothelial cells or pericytes was never observed. Instead SC were heavily embedded in the perivascular ECM, thus indicating that although SC share similar mechanisms to conquer the injured nervous systems, some differences exist in their mode of migration between CNS and PNS, and BV guidance and perivascular ECM seems to prevail for their migration in the CNS. The observed differences may result from different molecular and cellular environment existing between PNS and CNS, including different degrees of confinement in which SC are placed.

We previously demonstrated that SC avoid and are repelled by myelin(20, 58). Here we confirm these data and show that SC prevented to migrate directly through white matter, are somehow forced to migrate on BV. We showed previously that MAG, a CNS myelin component that prevents axonal regeneration in the CNS, is also inhibitory to SC migration and survival(20). While MAG accounted only partially for the repulsive effect of myelin to SC, our data in the developmentally arrested SC (Krox20^null^) and those on SC expressing low levels of Krox20(39) hint that the better blending with myelin observed in immature stages of the PNS-linage(37, 38) could be due to the lower expression of Eph receptor profile. We identified EphrinB3 as another myelin component negatively regulating SC in contact with CNS myelin. Like MAG, EphrinB3/EphA4 receptor signaling has been implicated in axon pathfinding(27). This suggests that myelin components exerting their inhibitory effect on axons are not exclusively directed against axons, but extend their inhibition to other neural components such as myelin-competent cells, preventing differentiation of oligodendrocyte progenitors into mature oligodendrocytes(59) as well as SC survival, and migration.

We first proved *in vitro*, that SC-bound EphrinB3 is able to activate by phosphorylation both EphA4 and EphB1, as well as to bind EphB6, which can be also transphosphorylated EphB1(43). Once bound to these receptors, EphrinB3 impairs the adhesion of these cells to myelin proteins, diminishing their process extension onto their substrate. Our *in vitro* data indicate that interfering with both EphB6 and EphA4 do not show additional improvement of this myelin-SC repulsion compared to single interference. Eph receptor signaling is not as straightforward as one receptor binding to one ligand. To trigger efficient activation, Eph receptors must cluster, homotypically (one subtype of Eph receptor) or heterotypically (involving the oligomerization of different Eph receptor subtypes)(41). This lack of synergic effect suggests that the mechanism of Eph/ephrin activation in SC might be mediated by heterotypic recruitment independent of the initial receptor activation.

Ephrin signaling includes not only induced repulsion but also modulates expression of adhesion molecules(29-31). We show that myelin-associated EphrinB3 modulates SC adhesion and migration to ECM, in particular FN, and enhanced integrinβ1 expression, overruling SC inhibition by myelin *in vitro* and promoting their migration along BV to reach the demyelinated lesions *in vivo*. This event occurs in correlation with increased FN expression, among other ECM molecules, during BV remodeling in response to demyelination, suggesting that the increased expression of ECM molecules by BV favours SC-BV interaction and subsequent migration along the vasculature *in vivo*.

Despite the present implication of EphA4 and EphB6 receptors in SC response to myelin, and their contribution to SC migration along CNS BV, the involvement of other Eph receptors in SC migration within the CNS should not be disregarded. To mention, EphB2, also expressed by SC and able to bind myelin-associated EphrinB3, mediates SC-SC interaction through N-cadherin re-localization to organize SC chain migration in the PNS(29). In fact, we observed SC in chains forming bridges going from one BV to another (Fig. 1C_2_ and F, Movie S1), suggesting the possible implication of EphB2 in these events.

In addition, EphA4 is involved in SC-astrocyte repulsion(28). Although the interactions between SC and astrocytes has not been explored in this study, the localization of the grafted SC in the vascular unit between the blood vessel wall and the astrocyte feet, suggests that the repulsion exerted by astrocytes can help confining SC to the perivascular space, and contribute to the mechanism of SC migration along BV. Interfering with EphA4 in SC could have altered this interaction and further allowed the grafted cells to escape from their perivascular path and mingle more with the surrounded white matter parenchyma.

In conclusion, we used multiple ex-vivo, in vivo and in vitro approaches to highlight a novel mechanism of guidance and migration of SC during the early events of CNS repair. We also provide strong evidences that the Eph/ephrin family regulates the complex interactions existing between SC myelin and blood vessels. SC encountering myelin-associated EphrinB3, retract their processes failing to mingle with white matter, and adhere preferentially to BV via activation of Integrinβ1. This dual effect, repulsing SC from CNS myelin and enhancing their attraction to basal lamina, directs their migration along CNS vasculature towards the lesion (Fig.9). Lesions of white matter undergoing the formation and/or reshaping of the vasculature with increased expression of ECM adhesion molecules, in particular FN, further triggers SC mobilization throughout the lesion. While SC invasion of the CNS is not restricted to demyelinating diseases, future studies should indicate whether such mechanisms are of relevance for other clinical pathologies such as trauma as well as genetic and acquired myelinopathies.

**Figure 9.**
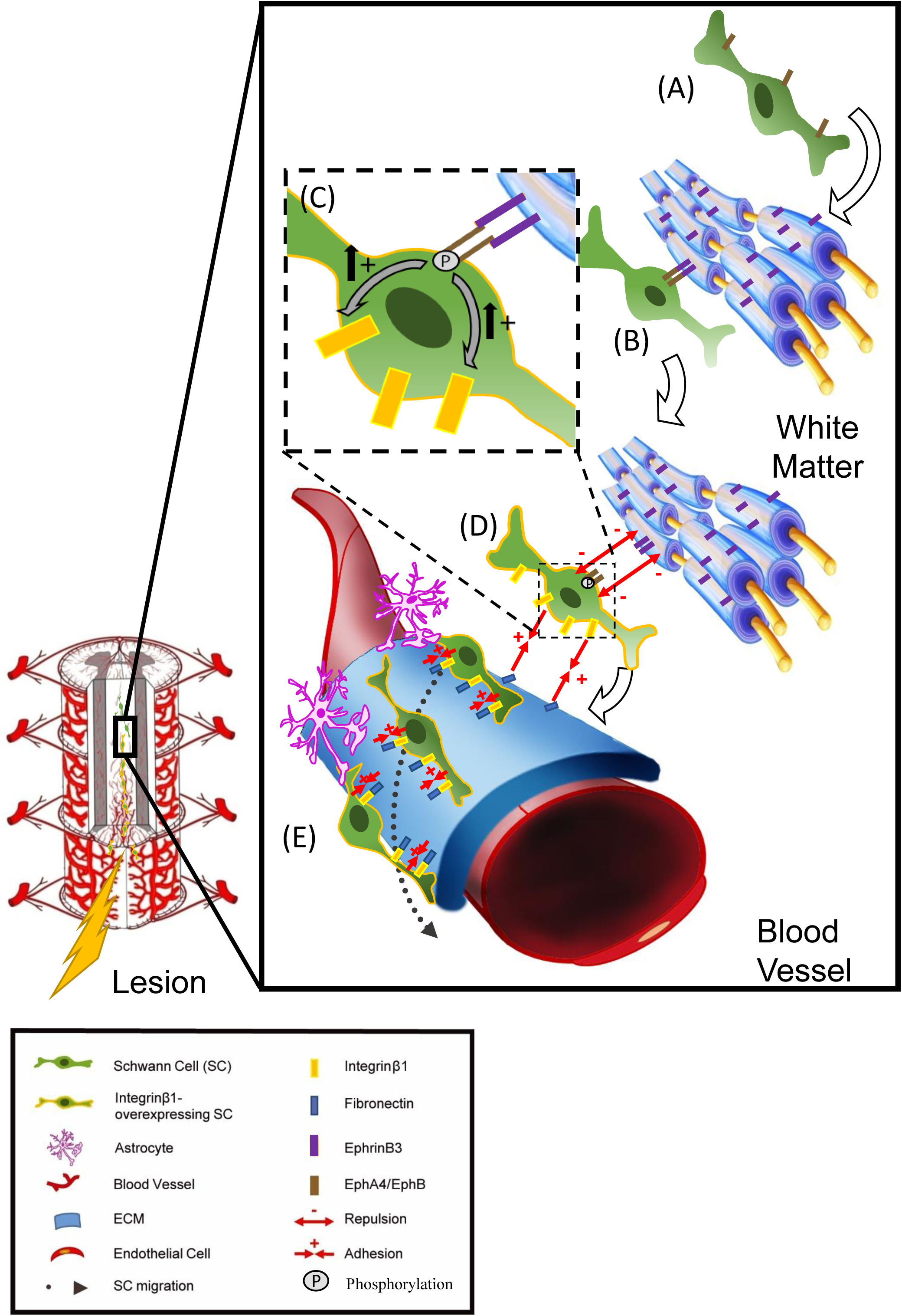
Model of the mechanism of guidance and migration of SC after CNS demyelination. (**A**) SC encountering CNS white matter (**B**) are activated by the myelin-associated EphrinB3 through EphB6-and EphA4-SC receptors. (**C, D**) The activation by phosphorylation of these receptors impairs SC adhesion to white matter and increases SC expression of Integrinβ1 promoting their adhesion to BV extracellular matrix (**E**). Lesions of white matter undergo the formation and/or remodeling of BV which increase expression of ECM adhesion molecules, such as FN, and further facilitate SC mobilization towards the lesion.

## METHODS

### Animals

Eight week old C57Bl/6JR female mice were purchased from Janvier Labs (Rodent Research Models & Associated Services). Actin-green fluorescent protein (GFP) transgenic, *Krox20*^*Cre/+*^ *R26R*^*YFP/+*^, and *Krox20*^*Cre/flox*^,*R26*^*mT/+*^ mice were previously characterized(3, 34), and maintained at ICM and IBENS animal facilities. Animal experiments were performed according to European Community regulations, ICM and INSERM ethical committee (authorization 75-348; 20/04/2005) and were approved by the local Darwin ethical committee.

### Schwann cell isolation and purification

Sciatic nerves of wild-type and Tg mice were isolated at postnatal day 15 and purification procedure was adapted from the previously described protocol(35). Briefly, enzymatic dissociation was performed by incubation with trypsin 0.025% and collagenase (420 U/ml) 10min at 37°C, followed by mechanical dissociation through different needle gages. After ending dissociation with fetal calf serum (FCS), SC were seeded in FN-coated flasks, and expanded in Dulbecco’s modified Eagles medium, containing 10% heat-inactivated FCS serum, penicillin (100 mg/ml), streptomycin (100 U/ml), human recombinant Neu-differentiation factor ß (hrNDFß) (125 ng/ml), insulin (10 µg/ml) and forskolin (2µg/ml).SC were purified by differential adhesion(60), obtaining a purity of 90%, and passaged no more than three times before use. Purification was controlled by immunocytochemistry for p75.For adhesion, migration and blocking receptor assays, SC were maintained in Sato serum-free medium(61) supplemented with hrNDFB (125 ng/ml), and forskolin (2 µg/ml).

### iDISCO whole-mount immunofluorescence and imaging

Spinal cords were processed as described in the iDISCO protocol(62), including modifications described in the updated online protocol (https://idisco.info, December 2016). The primary antibody used was rabbit anti-RFP (1:1000, Rockland). Secondary antibodies used were donkey anti-rabbit Cy3 (1:800, Jackson Immunoresearch) and donkey anti-mouse IgG Cy5 (1:800, Jackson Immunoresearch) for intravascular staining. The cleared samples were imaged with a light sheet microscope (Ultramicroscope II; LaVision Biotec).

### RNA transcriptome analysis

Library preparation and Illumina sequencing were performed at the EBENS genomic core facility. Briefly, (polyA+) mRNAs were purified from 250 ng of total RNA using oligo(dT). Libraries were prepared using the strand specific RNA-Seq library preparation TruSeq Stranded mRNA kit (Illumina). Libraries were multiplexed by 6 on 1 high-output flow cells. A 75 bp read sequencing was performed on a NextSeq 500 device (Illumina). A mean of 94 ± 9,5 million passing Illumina quality filter reads was obtained for each of the 6 samples.

The analyses were performed using the Eoulsan pipeline(63), including read filtering, mapping, alignment filtering, read quantification, normalization and differential analysis. Before mapping, poly N read tails were trimmed, reads ≤40 bases were removed, and reads with quality mean ≤30 were discarded. Reads were then aligned against the Mus musculus genome from Ensembl version 84 using STAR(64). Alignments from reads matching more than once on the reference genome were removed using Java version of SamTools(65). To compute gene expression, Mus musculus GFF3 genome annotation version 84 from Ensembl database was used. All overlapping regions between alignments and referenced gene were counted using HTSeq-count 0.5.3(66). The sample counts were normalized using DESeq 1.8.3(67). Statistical treatments and differential analyses were also performed using DESeq 1.8.3.

### Data availability

The RNASeq gene expression data and raw fastq files are available on the GEO repository (www.ncbi.nlm.nih.gov/geo/) under accession number: GSE107401 (accession password: mlkfkwoezxujvsn).

### Myelin Protein Extract isolation

Myelin was purified by sucrose gradient centrifugation(68). Cerebral hemispheres of adult mice (3 months-old)were homogenized (on ice) in 0.35 M sucrose and 5 mM EGTA and the suspension was overlaid onto an equivalent volume of 0.85 M sucrose and 5 mM EGTA and centrifuged at 100000 x g at 4°C for 20 min. The myelin-containing fraction at the interface was collected, diluted three-fold in distilled water, and centrifuged at 100000 x g at 4°C for 30 min. After washing with distilled water, the isolated myelin pellet was re-suspended in 20 mM Tris-HCl, aliquoted, and stored at –20°C.

### Pre-clustering of recombinant EphrinB3-Fc

Mouse EphrinB3-Fc fragments and human Fc were purchased by R&D Systems. The soluble forms of EphrinB3-Fc, and its control Fc, have low effect on receptor activation(69), therefore they were mixed with anti-mouse Fc-IgG and anti-human Fc-IgG (Alexa 555) respectively, (ratio = 1:5), and incubated for 1 h at 37°C prior to addition to SC(70).

### Adhesion and Spreading Assays

*In vitro*. Adhesion and spreading assays were performed in 24 well dishes. Silicon strips on coverslips were used to separate two coated areas of each coverslips(71). Surfaces were coated overnight at 37°C with recombinant EphrinB3-Fc fusion at 10 µg/mL and Fc equimolar (as control) on each side respectively; or myelin extract (100 µg/mL) and PBS buffer, as control. Before cell seeding, strips were removed and coverslips were washed carefully with PBS. 10^5^ SC were seeded in serum-free Sato medium to avoid proliferation, and allowed to adhere for 3h. Data were always expressed as ratio respect to the intra-coverslip control(24).

### Spinal cord frozen sections

Cryostat longitudinal sections(20µm thick)of snap frozen spinal cords were allowed to adhere to glass slides and were thawed at room temperature just before use. Then, 50.000 SC/sections were seeded and allowed to adhere overnight (16h) before fixation in 4% paraformaldehyde (5 min), immunostaining for Glut1and mounting with fluoromount.

### Survival assay

GFP^+^SC were seeded on uncoated glass coverslips in normal medium. After overnight adhesion, medium was changed adding Sato serum-free medium supplemented with clustered-EphrinB3 at 10 µg/mL and Fc equimolar (as control); or myelin extract (100 µg/mL) and PBS (as control). SC were incubated for 3 h or 24 h as specified in each experiment. After fixation in 4% paraformaldehyde (5 min), SC were immuno-stained for caspase 3 adding Hoechst dye to visualize all nuclei, and coverslips mounted with fluoromount.

### Migration assay.SC were re-suspended at 3·10^6^ cells/ml in Sato medium containing 0.8% low melting point agarose (Sigma)

1.5 µL-drops of this suspension were applied to the center of FN +EphrinB3, or FN +Fc-coated glass coverslips, which were placed at 4°C for 1 min to allow the agarose to solidify. The cooled drop was covered with Sato medium with hrNDFß(125 ng/ml) and forskolin (2 µg/ml) and placed up to 6 h at 37 °C into the incubating chamber of a video-microscopy (ZEISS).

### SC receptor blocking assay

EphA4 and EphB6 receptors, or Integrinβ1 in SC were neutralized by incubation with anti-EphA4 (1.2 µg/10.000 cells, R&D, AF641), anti-EphB6 (1.2 µg/10.000 cells, Santa Cruz Biotechnology, sc-7282), anti-integrinβ1 (0.6 µg/10.000 cells, MA2910, Thermo Fisher Scientific) antibodies or IgG (as control) in Sato medium for 1 h at 37°C prior to cell seeding or transplantation.

### Immuno-staining

Cultured SC were fixed for 5 min in 4% paraformaldehyde prior immuno-staining and mice were sacrificed by trans-cardiac perfusion of PBS followed by cold 4% paraformaldehyde, and post-fixed in the same fixative for 1 hour. Spinal cords were cryo-protected by immersion in 20% sucrose solution overnight, embedded in cryomatrix (Thermo Scientific), and frozen in cold isopentane at –60°C. Finally, they were sectioned with a cryostat at 12 µm (Leica Microsystems). Both cells and sections were washed, blocked in 5% BSA for 40 min and incubated with the primary antibodies. While cells were incubated 1h at room temperature, slides were incubated overnight at 4°C. For MOG staining, sections were incubated with absolute ethanol for 10 min followed by primary antibody, and then washed profusely. Primary antibodies were as follows: anti-EphA4 (1:50, R&D, AF641); anti-EphA4-Tyr(602) (1:50, ECM Biosciences, EP2731); anti-EphB6 (1:50, SAB4503476, Sigma); anti-EphB1 (1:50, SAB4500776, Sigma; anti-Eph receptor B1+Eph receptor B2 (phospho Y594) (1:50, ab61791, Abcam); anti-Ki67 (1:100, 556003, BD Biosciences); anti-cleaved caspase3 (1:500, Cell Signalling, #9661S); anti-GFP (1:400, Aves, GFP-1020); anti-MOG (1:20, mouse IgG1 hybridoma, clone C18C5; provided by C. Linnington, University of Glasgow, Glasgow, United Kingdom); anti-MBP (1:50, ab7349, Sigma), anti-Glut1 (1:100, 07-1401, Merck Millipore; and 1:400, MABS132, Sigma); anti-Fibronectin (1:600, F6140, Sigma); anti-CD31 (1:200, 553370, BD Pharmigen); anti-NF200 (1:200, N4142,Sigma), anti-p75 (1:100, Ozyme, 8238S), anti-CD13 (1:50, BioRad, MCA2183), anti-CD68 (1:400, BioRad, MCA1957), anti-CD11b (1:400, BioRad, MCA74G), anti-F8/40 (1:100, BioRad, MCA497R), anti-Collagen 4(1:400, ab19808, Abcam), anti-Olig2 (1:300, MABN50, Millipore) and anti-sox10 (1:50, AF2864, R&D System).Next, cells or sections were washed and incubated with secondary antibodies and Hoechst dye for 1h at room temperature. The excess secondary antibody was removed by several PBS washes, and coverslips/slides were mounted using fluoromount.

### Electron microscopy

For electron microscopy, mice were perfused with PBS followed by 4% paraformaldehyde/2.5% glutaraldehyde (Electron Microscopy Science) in PBS for 45 minutes. Dissected spinal cords were post-fixed with the same solution for 2 hours, then, sectioned into 60µm slices with a vibratome and washed twice with PBS before enzyme immuno-labeling revealed by DAB/oxygen peroxyde. For DAB revelation, endogenous peroxidase was inhibited with a methanol/oxygen peroxide incubation, washed and blocked by 5% BSA for 1h. Sections were incubated with anti-GFP overnight at 4°C, then washed and incubated with a secondary biotinylated antibody for 2h at room temperature. After several washes with PB 0.1M sections were incubated with the ABC kit (VECTASTAIN® ABC-HRP Kit, Vector Lab) containing peroxidase-anti-peroxidase for 40min followed by a DAB/ oxygen peroxide mix before stopping the reaction with distilled water. Samples were fixed in 2% osmium tetroxide (Sigma-Aldrich) 30min, washed gently and incubated with 5% uranyl acetate for 30min in the dark. After dehydration, samples were embedded in Epon resin 812. Ultra-thin sections (80 nm) were examined with a HITACHI 120 kV HT-7700 electron microscope.

### Demyelinating lesions and grafts

Wild-type mice were anaesthetized with a ketamine/xylazine mixture. Demyelination was induced by stereotaxic injection of LPC (1%, 0.5 µl) in PBS. LPC or PBS (in control animals) was injected into the dorsal funiculus of the spinal cord at the level of T8–T9 in the dorsal column white matter using a glass micropipette. SC (10^5^/ 2µL) were injected the same day two vertebrate caudally (4mm) in the same tract.

### Western blotting

SC (6·10^6^ cells/well) were lysed in RIPA buffer with Complete® and Phosphostop® inhibitor and analyzed by electrophoresis in an SDS 4–20% MINI PROTEAN TGX gel. After electrophoresis, proteins were transferred electrophoretically to polyvinylidene difluoride membranes and probed with the following antibodies:anti-EphB6 (1:500, SAB4503476, Sigma), anti-EphB1 (1:500, SAB4500776, Sigma), anti-EphA4 (4µg/mL, 37-1600, ThermoFisher), anti-p-EphB1+2 (1:300, ab61791, Abcam), anti-p-EphA4 (1:300, EP2731, ECM Biosciences), anti-Integrinβ1 (1:500, 550531, BD Pharmingen), anti-EphrinB3 (1:250, 1µg/mL, AF395 R&D systems), anti-MBP (1:1000, ab980, Millipore), anti-GAPDH (1:5000, MAB374, Millipore) and anti-Actin (1:50000, A2228, Sigma). Peroxidase-conjugated anti-rabbit, anti-goat or anti-mouse IgG secondary antibodies (Jackson Immuno Research) were used at a dilution of 1:5000, 1:10000 and 1:20000, respectively, and anti-Rat biotinylated (1:100, Vector Labs) followed by peroxidase-conjugated streptavidin. Protein bands were visualized by chemoluminescence (ECL BioRad). Intensity of the bands was quantified with ImageJ.

### Neutralization of EphrinB3 epitopes in myelin extracts

EphrinB3 epitopes in myelin extract proteins were neutralized by incubation with anti-EphrinB3 antibodies (AF395, R&D system and sc-271328, Santa Cruz Biotechnology; ratio: 1:1) for 2 h at room temperature prior to the addition to the cells(26).

### Spinal cord live imaging

LPC lesion followed by GFP^+^SC engraftment was performed in 60 days-old animals and mice terminally anesthetized. Rhodamine labeled BSL I (Vector Labs RL-1102) at 2mg/ml was injected in the beating heart to label BV. After 5 minutes allowing dye circulation, spinal cords were dissected in ice-cold HBSS solution supplemented with 6,4 mg/mL d-(+)-glucose and bubbled for 30 min with bubbled with 95% O_2_/5% CO_2_. Spinal cord segments including lesion and graft sites were laid onto Millicell-CM slice culture inserts (Millipore) over culture medium (50% DMEM+Glutamax, 25% HBSS, 25% heat-inactivated horse serum, 5 mg/mL d-(+)-glucose, 20mM Hepes, penicillin (100 mg/ml), streptomycin (100 U/ml), human recombinant neu-differentiation factor ß (hrNDFß) (125 ng/ml), and forskolin (2 µg/ml)in a glass bottom plates, and then placed into an inverted Leica SP8X confocal with an on-stage incubator (while streaming 95% O_2_, 5% CO_2_ into the chamber). Spinal cords were imaged using a 25x immersion objective at intervals of 15 min during 12 hours with intermittent repositioning of the focal planes. Maximum intensity projections of the collected stacks (∼60μm at 2μm step size) were compiled in FIJI program.

### Quantification

#### Lesion

The lesion area was identified by immune detection of GFAP combined with Hoechst (H^+^) labeled nuclei to reveal astrocyte reactivity and hyper-cellularity respectively.

#### SC adhesion and spreading in vitro

SC adhesion on different surfaces was quantified as the ratio of the number of adhered GFP^+^ on myelin-or EphrinB3-coated areas, over those adhered to uncoated or Fc-coated area within the same coverslip. All coated areas were of equal size. Schwann cell spreading was evaluated by quantifying the ratio of GFP^+^SC not able to expand their processes out of the total on myelin-or EphrinB3-coated areas over those on non-coated or Fc-coated.

#### SC spreading and adhesion to BV ex vivo

SC spreading on frozen spinal cord sections was evaluated by quantifying the GFP^+^ area per cell identified by Hoechst^+^ nuclei. Adhesion on these sections was evaluated as the number of cells in contact with collagen 4-positive blood vessels.

#### SC extent of migration in vitro

SC migration was quantified by measuring the number of SC outside the 1.5 µL agarose drop and the maximum extent of their migration from the edge drop.

#### SC velocity in vitro

SC speed of migration was quantified by manual cell tracking plugging of FIJI program, calibrating pixel size and duration of time-lapse of each frame.

#### Size of lesion and graft area

Lesion and grafted cells within the dorsal funiculus were quantified by delimiting Hoechst+ nuclei hyperdensity and GFAP-positive area per 12µm-section. Lesion and graft areas were quantified by ImageJ 1.49s. For each animal, at least three serial sections with 60 µm intervals were quantified.

#### Extent of SC migration in vivo

SC migration within the dorsal funiculus was quantified on longitudinal sections evaluating the distance between the graft injection site and the most proximal GFP^+^ cell to the lesion (LPC injection site) in each animal from different groups.

#### BV expansion

BV were identified by immunoreactivity to Glut1, a marker restricted to micro-vessels with blood-tissue barrier function. BV expansion was quantified as the Glut1^+^area over a cut-off threshold of staining using ImageJ 1.49s.Glut1^+^area in controls was averaged and every animal value of both groups was expressed as a ratio over this control mean. Endothelial cell proliferation was assessed taking into account only Glut1^+^ cells containing a Ki67^+^ nucleus clearly embedded within their cytoplasm, and excluding Ki67^+^ nuclei, closely apposed.

#### SC-BV association

SC-BV association was quantified in the intermediate zone, at1mm from the graft edge in the direction of the lesion or in the lesion site. GFP^+^/Hoechst^+^ SC, with complete or more than half-cell size in contact with Glut1^+^ endothelial cells were counted as “closely associated cells” while those with only “tip” contacts or no contacts were considered as “not associated cells”. Data are expressed as the percentage of total counted cells in both groups.

#### SC-Axon alignment

SC-BV alignment in the lesion site was quantified. GFP^+^/Hoechst^+^ SC, with complete or more than half-cell in parallel orientation and aligned closely to NF200^+^ axons (excluding alignment to BV), were counted as “closely associated cells” while those with only “tip” contacts or no contacts were considered as “not associated cells”. Data are expressed as the percentage of total counted cells in both groups.

### Statistics

The sample size calculation was performed by the resource equation method, trying to minimize the sample size to follow the ARRIVE guidelines for reporting animal research. Each n represents one animal or SC sample in the experiment. The grafting experiments were repeated at least three times with different set of animals each. For the *in vitro* analysis, experiments were performed at least three times with SC obtained from different dissections and dissociations. Statistical analysis was carried out using GraphPad Prism 6 software. All values were expressed as mean ± SD. Normality in the variable distributions was assessed by the D’Agostino&Pearson omnibus test and Grubbs’ test was used to detect and exclude possible outliers. When Normality test was passed, means were compared by two-tailed Student’s t test. When one or both groups did not follow a normal distribution, means were compared by two-tailed Mann-Whitney U test. When different independent groups were compared, we performed a one-way ANOVA plus Tukey’s multiple comparison tests. One sample t-test was used to compare values to the hypothetical mean: 1 for ratios and 100 for percentages. Repeated-measure ANOVA was used to analyze the difference along time of a certain parameter. P-values lower than 0.05 were used as a cut-off for statistical significance.

## AUTHOR CONTRIBUTIONS

B.G-D. coordinated and designed the study, performed most of the experimental work, analyzed and interpreted the results, and wrote the manuscript with critical input from all authors; C.B. performed the *ex vivo* immunohistochemistry experiments; F.C. carried out gene profiling and analysis; G.G. performed iDisco and LightSheet analysis; C.D. assisted with animal experiments; V.Z., P.T., and P.C. critically reviewed the manuscript; A.B-V. conceived the project, handled the funding, supervised and interpreted the results, wrote and edited the manuscript. All of the authors approved the manuscript.

## ACKNOWLEDGMENTS

The authors thank the ICM and IBENS animal, genomic and cellular imaging core facilities, and the ICM histology core facility for technical assistance. The IBENS Imaging facility received the support of grants from the “Région Ile-de-France” (NERF N°2011-45), the “Fondation pour la Recherche Médicale” (N° DGE 20111123023), and the “Fédération pour la Recherche sur le Cerveau - Rotary International France”(2011). The ICM core facility received funding from the ICM Foundation.

Funding was provided by grants from the National Multiple Sclerosis Society (NMSS), to A.B. and P.C.(RG 5088-A-1); INSERM, CNRS, ARSEP, and the program “Investissements d’Avenir” (ANR-10-IAIHU-06 and ANR-11-INBS-0011– NeurATRIS) to AB; INSERM, CNRS, IBENS, the program “Investissements d’Avenir” (ANR-10-IAIHU-06 and ANR-11-INBS-0011–NeurATRIS) to P.C; the France Génomique national infrastructure, funded as part of the “Investissements d’Avenir” program (contract ANR-10-INBS-09) to P.T; Junta de Andalucía and the European Commission under the Seventh Framework Programme of the European Union (agreement Num. 291730, contract TAHUB-II-107) to B.G-D.

