## Supplementary figures and note for "Blood vessels guide Schwann cell migration in the adult demyelinated CNS through Eph/ephrin signaling"

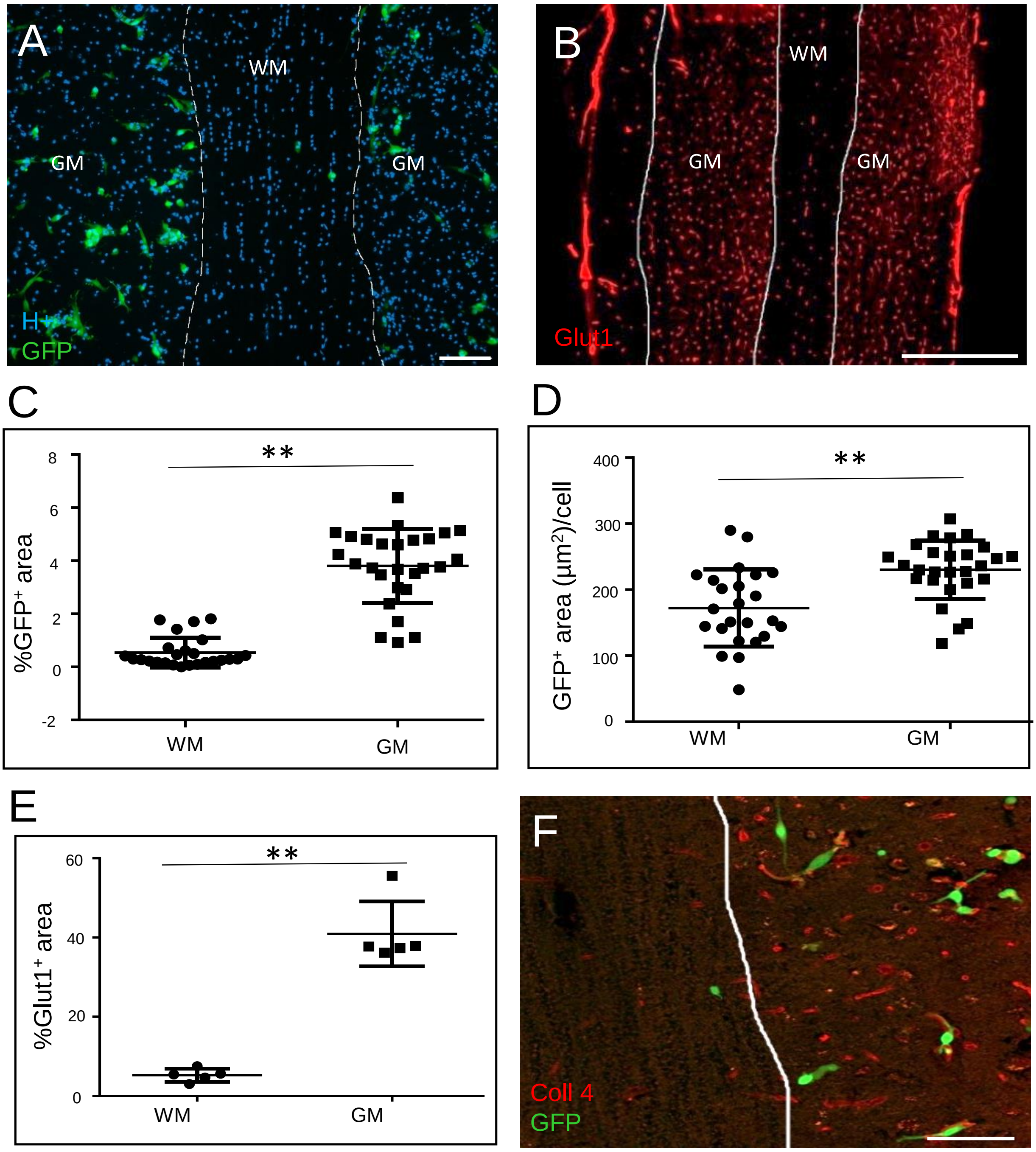


**Figure S1. Schwann cells adhere less on CNS white matter, but associate preferentially to blood vessels *ex vivo*.** (**A**) GFP$^{+}$SC seeded *ex vivo* on frozen spinal cord sections, adhere less over white matter (WM) than over grey matter (GM), scale bar 100 µm. (**B**) Glut1$^{+}$ BV are more abundant in GM than WM, scale bar 500 µm. (**C**) Evaluation of GFP$^{+}$ area confirms SC greater affinity for GM (n=27) over WM (n=21) (two-tailed Mann Whitney test p<0.001). (**D**) Evaluation of GFP$^{+}$ area per cell shows an increase of SC spreading in GM (n=27) over WM (n=24) (two tailed t-test p=0.0002). In **C** and **D** (n) represents random fields of 3 different experiments repeated independently twice, mean±SD. (**E**) Quantification of Glut1$^{+}$ area in GM over WM (two-tailed Mann Whitney test p=0.0079) (n=5 different experiments repeated independently three times, mean±SD). **(F)** Co-detection of collagen 4 (Coll 4,) and GFP showed that SC adhere preferentially to BV, scale bar 100 µm. * means p<0.05, ** means p<0.01

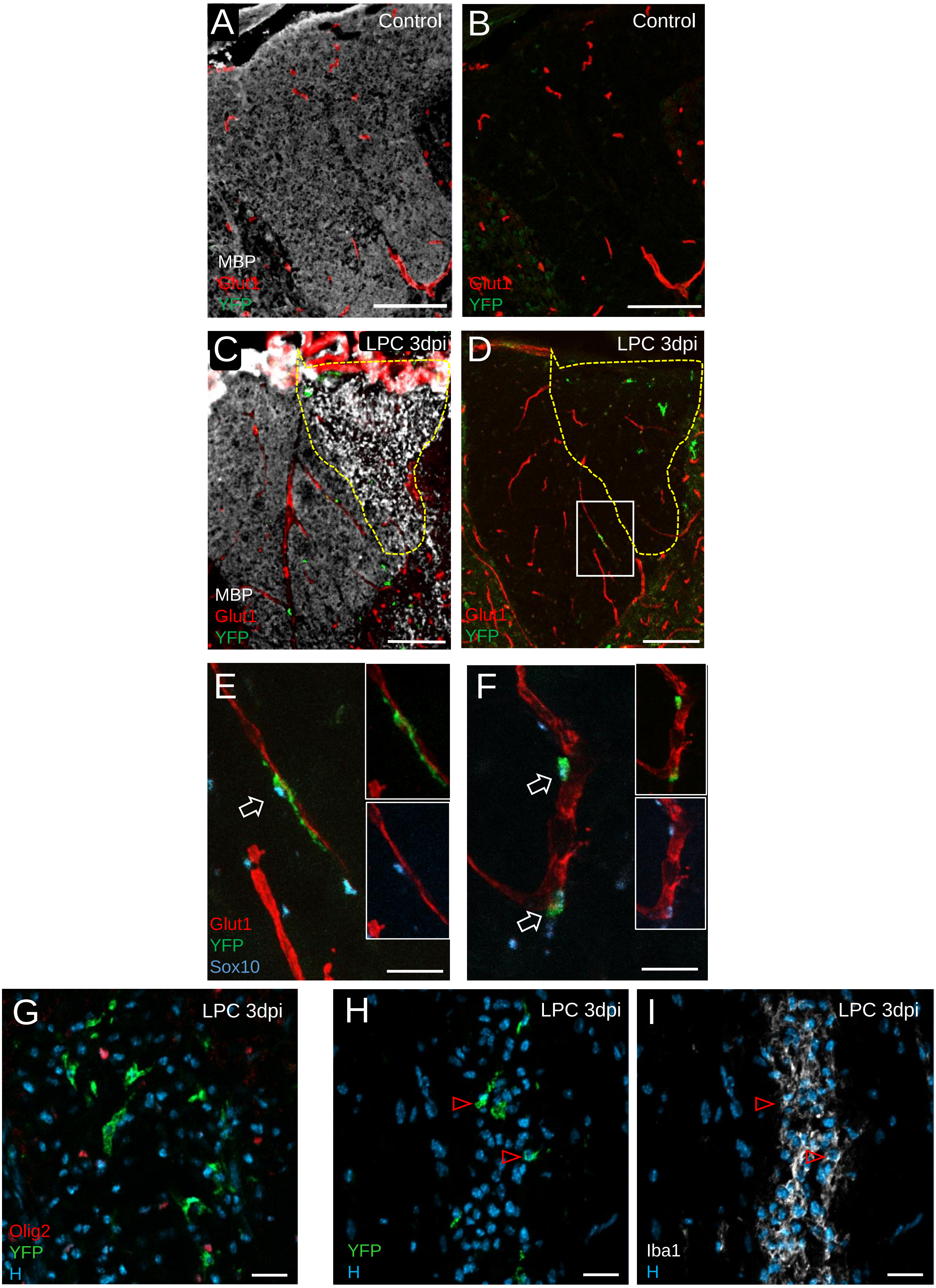


**Figure S2. Perivascular migration of endogenous SC in response to demyelination.** (**A,B**) MBP, Glut1 and YFP immunostainings of control *Krox20*$^{Cre/+}$ *R26R*$^{YFP/+}$spinal cord sections show the absence of YFP$^{+}$SC in spinal cord without lesion. (**C,D**) General view of YFP$^{+}$SC on Glut1$^{+}$ BV near the lesion at 3dpi. (**E,F**) Examples of YFP$^{+}$ SC associated with Glut1$^{+}$ BV and expressing Sox10; insets show separate colors for YFP and Sox10; E is an enlargement of the boxed area in **D**. (**G**) YFP$^{+}$ cells do not co-label with the oligodendroglial marker Olig2. (**H,I**) YFP$^{+}$ cells (green) express sometimes (red arrows) the microglial marker Iba1 (white). Scale bar: in A-D 100 µm, in E-F:20 µm.

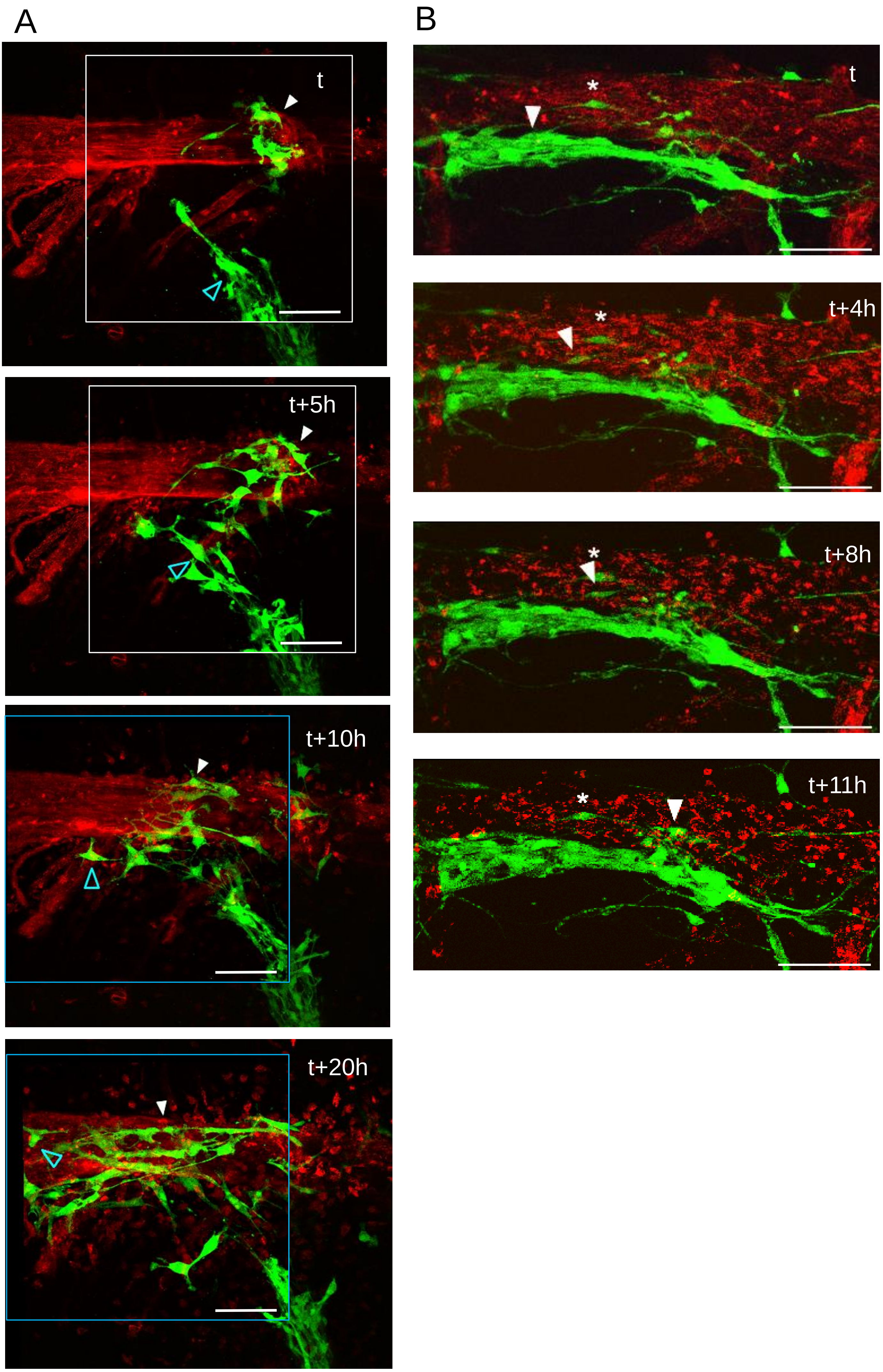


**Figure S3. Time-lapse imaging of SC movements on blood vessels.** (**A**) Grafted GFP$^+$SC (green) in the spinal cord reach rhodamine-lectin labeled BV (red). The white arrowhead follows a GFP$^+$SC gliding along a large BV, the blue arrowhead follows a GFP+SC jumping from one BV branch to another (movie S1). (**B**) GFP$^+$SC tend to migrate in chain on BV, but some (white arrowhead) escape and move on the outer BV surface; the star identifies a SC that is less motile (movie S2). Scale bar 100 µm.

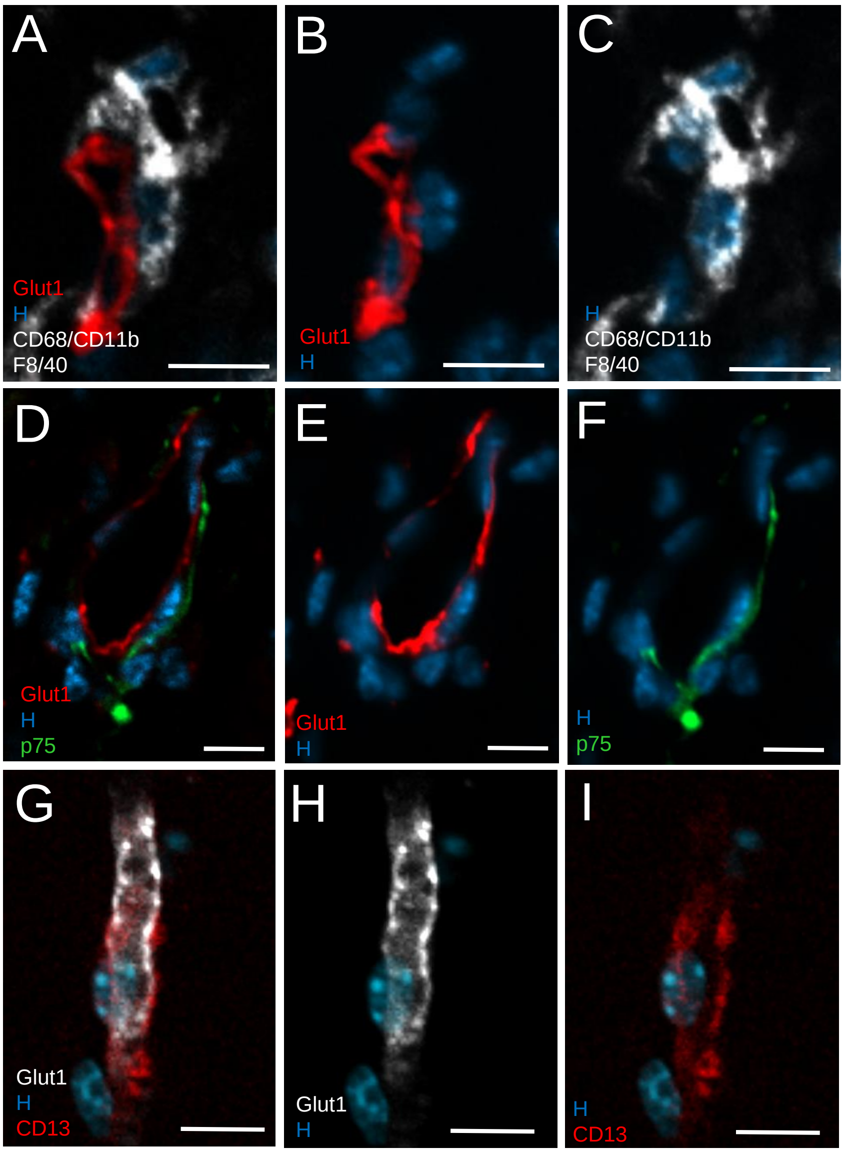


**Figure S4. Glut1 specifically labels endothelial cells.** (**A-F**) Absence of co-labeling of Glut1 and (**A-C**) the microglial/macrophages markers CD68, CD11b and F8/40, (**B-F**) the SC marker, p75, (**G-I**) the pericyte marker, CD13. Scale bar 10 µm.

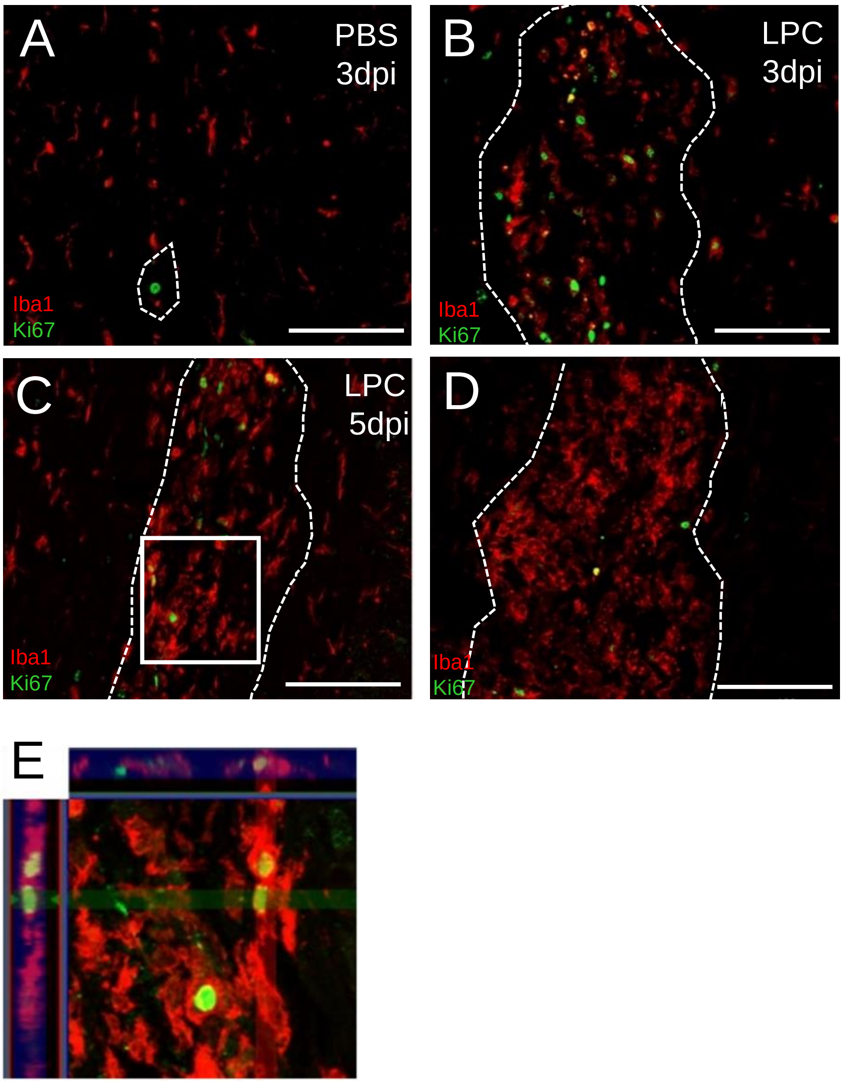


**Figure S5. Proliferative response of microglial cells to LPC-induced demyelination of the adult spinal cord.** (**A-D**) Immunostaining for Ki67 (green) in combination with Iba1 (red) identifies proliferating cells after PBS injection at 3dpi (**A**), and after LPC injection at 3 (**B**), 5 (**C**) and 7 (**D**) dpi. Most Ki67$^{+}$ nuclei present in the lesion (outlined by dotted lines) were Iba1$^{+}$ microglial cells. Ki67$^{+}$/Iba1$^{+}$ cell numbers decrease with time and are nearly undetectable in PBS injected animals (**A**). (**E**) Orthogonal projection of the boxed area in **C**, confirms that Ki67-positive cells are Iba1$^{+}$. Scale bar 100 µm

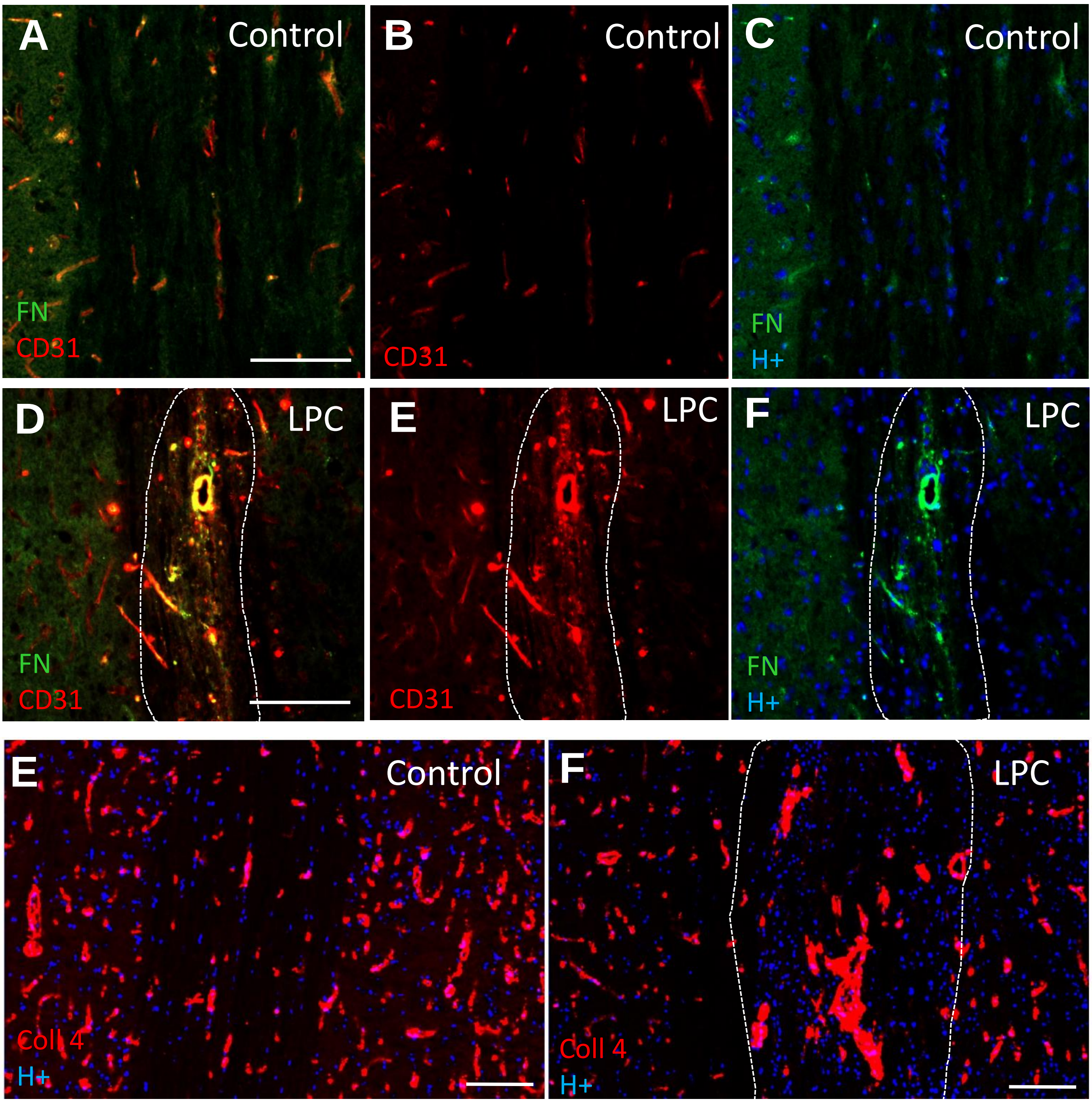


**Figure S6. Dynamics of ECM expression in response to LPC induced demyelination of the spinal cord.** Immunodetection of FN (**A,C,D,F**) and CD31 (**A,B,D,E**) indicates that FN expression by BV increases significantly in LPC lesions (**D-F**) compared to control (**A-C**). (**E,F**) Collagen 4 expression is also enhanced around BV (**F**). Dashed lines identifies the lesion borders. Scale bar 100 µm.

| *Gene* | *WT* | *Mut_krox20* | ***log2FoldChange Mut vs WT*** | *pval* |
|---|---|---|---|---|
| Eph receptor A4 | 755,7122 | 226,0449 | **-1,7412** | 3,4343E-12 |
| Eph receptor B1 | 274,7597 | 64,7978 | **-2,0842** | 4,7769E-06 |
| Eph receptor B2 | 1909,5945 | 2664,8813 | **0,4808** | 0,01258419 |
| Eph receptor B3 | 2194,4845 | 2511,7291 | **0,1948** | 0,2936923 |
| Eph receptor B4 | 3541,5449 | 2695,7100 | **-0,3937** | 0,07857419 |
| Eph receptor B6 | 7149,8441 | 921,0603 | **-2,9565** | 1,7032E-39 |

**Figure S7. RNA sequencing analysis.** Comparison of both Krox20$^{null}$ and wild-type SC expression profiles shows reduced levels in EphA4, EphB1 and EphB6 transcripts of mutant SC, n=3.

**Supplementary note 1.**

Specificity of endothelial cell proliferation was assessed by co-immunolabeling Glut1 with different lineage markers. Glut1$^+$ cells did not express the pan microglial/macrophage markers (CD68, CD11b,F8/40) (Fig. S4A,C), the SC precursor and non myelinating SC marker p75 (Fig. S4D,F), the pericyte marker CD13 (Fig. S4G,I) nor the SC/astrocyte marker GFAP (Fig. 1E), thus confirming the specificity of Glut1 for endothelial cells. Moreover confocal imaging excluded potential overlap with proliferating perivascular cell types. Endothelial cell proliferation within the lesion was mild over the total number of Ki67$^+$ cells (3±1% of the total Ki67$^+$ are Glut1$^+$), the latter corresponding mainly to proliferation of microglial cells labeled with Iba1 (Fig. S5) and possibly few glial cells (astrocytes, OPC, SC) present in the lesion at this early time-point.
